# Extraordinary activation of CALB by alkylammonium ions: a new paradigm for activity enhancement of enzymes

**DOI:** 10.1101/2025.04.18.649601

**Authors:** Rangasamy Savitha, Ebin K. Baby, Gemma K. Kinsella, Kieran Nolan, Barry J. Ryan, Gary T. M. Henehan

## Abstract

*Candida antarctica* lipase B (CALB) is widely used in biocatalysis with applications in plastics degradation and chemical synthesis. CALB is activated by hydrophobic matrices and, enigmatically, shows striking activation in polar, choline-based, Deep Eutectic Solvents (DES). Herein, we show that CALB activation and stabilisation by TAAs is caused by binding to choline’s tetraalkylammonium (TAA) moiety. Several related TAA salts also caused CALB activation which was proportional to the hydrophobicity of their alkyl substituents. Remarkably, tetraoctylammonium bromide showed activation of ∼500% even at low micromolar levels. These TAA salts represent a new class of enzyme activator. Molecular modelling identified the alkylammonium binding location as a hydrophobic patch centred around Asp-145 of CALB. Binding at this site explains lipase activation in polar DES solvents and its relationship to other pathways of CALB activation.

Herein, we also demonstrate that CALB, like many lipases, is activated by calcium. Intriguingly, mixed soluble activator experiments showed that calcium and choline bind to different CALB sites, suggesting a two-site model for CALB activation.

These observations, along with previous findings, show that TAA activation is a widespread property of enzymes and constitutes a novel and potent means to enhance enzyme turnover and stability.

**Highlights:**

1. CALB is activated by choline
2. Several tetraalkylammonium salts cause activation of CALB
3. Hyperactivation of CALB (5-fold) by tetraoctylammonium ions occurs at low micromolar concentrations.
4. Two independent sites for CALB activation, by calcium and TAA ions, are identified
5. Activation at the choline binding site stabilises CALB while calcium binding destabilises the enzyme
6. A soluble activator is demonstrated, that can be used to probe the activation mechanism of CALB or other enzymes.

## 1 Introduction

Since the pioneering work of Klibanov, there have been steady advancements in the application of enzymes in organic synthesis.^1^ Enzyme biocatalysis is of ever-increasing interest to synthetic chemists due to a growing focus on green chemistry solutions.^2,3^ Despite the widespread use of enzymes in biocatalysis, there are still drawbacks to their use in many application areas due to limitations in enzyme activity, solvent tolerance and thermal stability.^4,5^ To address this deficit, extensive efforts, over many years, have been made to enhance enzyme performance.^2,4^ Early studies focussed on chemical modification to enhance enzyme properties^6^ while later years saw the adoption of site directed mutagenesis strategies.^4,5,7^ More recently, solvent engineering using neoteric reaction media such as ionic liquids or deep eutectic solvents (DES) has been explored.^8,9^ Based on these efforts to improve enzyme performance, it is clear that several important enzymes used in biocatalysis have significant “locked in” catalytic potential. For example, covalent modification of an alcohol dehydrogenase was reported to increase activity up to 20-fold.^10^ Often, relatively minor changes in key amino acids, by site directed mutagenesis or covalent modification, can lead to dramatic increases in turnover.^10–13^ Judicious choice of solvent can also dramatically improve enzyme turnover and examples of rate enhancement in DES abound. Thus, activation was reported for a cutinase, protease, carbohydrase, and other enzymes in neoteric media.^5,14^ Clearly, many enzymes are inherently capable of accessing activated states. Unlocking the latent catalytic potential of enzymes is of great commercial value since the enzyme is often the most expensive element of a biocatalytic process.^15,16^

Lipases, occupy a central position in biocatalysis and cover a broad range of applications. For instance, several lipases, and closely related cutinases, have been employed in plastics breakdown, transesterification reactions and in biodiesel synthesis.^17,18^ They are increasingly used for production of sustainable and biodegradable polyesters.^19^ Lipases also catalyse an unusually broad range of transformations in fine chemicals production including ester hydrolysis, ester synthesis (by reverse hydrolysis), olefin epoxidation and amide synthesis among others.^20^

Perhaps, the most striking case of enzyme activity enhancement is the interfacial activation reported for many lipases. The catalytic activity of lipases increases dramatically when they encounter a hydrophobic phase, typically an oil/water interface. This activation is due to the movement of an alpha helical loop of protein that covers the active site when it is in an aqueous environment. This “lid” limits substrate access to the active site. Contact with a lipid phase displaces this “lid”, opening the active site and greatly enhancing lipase turnover.^21,22^ While this activation holds true for several lipases, *Candida antarctica* Lipase B (CALB) (recently reclassified as *Pseudozyma antarctica* lipase B)^23^, exhibits weak or negligible interfacial activation due to its unique structural features. The lack of a prominent lid^24^ in CALB makes the catalytic site fully accessible to substrates in both aqueous and organic media. CALB stands out as a lipase with particularly favourable properties and occupies a central position in biocatalysis due to its inherent robustness and broad specificity.^20^

Despite its negligible interfacial activation, CALB was shown to be activated and stabilised when immobilised on hydrophobic supports. The benchmark biocatalyst, Novozyme 435, consists of CALB non-covalently immobilised on a cross-linked, macroporous, poly(methyl methacrylate) resin through hydrophobic interactions.^25^ CALB has a hydrophobic pocket near the active site which is thought to facilitate such interactions.^26^ In immobilised form, it displays an unusually high degree of tolerance to high temperatures, solvents and pH extremes and has been referred to as the “perfect enzyme”.^25^

Besides those mentioned above, two other significant strategies to activate lipases have been reported. The first, rests on the observation that several lipases become activated by surfactants.^27^ Of particular relevance to the present work, is the observation that cationic surfactants are unusually effective at lipase activity enhancement. Such activation is greater for surfactants with longer hydrophobic chains. For example, in an illuminating study, the cationic surfactant, cetyltrimethylammonium bromide (CTAB) showed activation of a lipase from *Thermomyces lanuginose*.^28^ It was also shown that longer chain length detergents were more effective at CALB activation, due to their increased hydrophobicity.

Secondly, it is worth noting that, as with many lipases, calcium activation is observed for CALB.^26,29^ Calcium ions are known to stabilise certain proteins through cross linking and some enzymes have specific binding sites for this ion.^30–34^ However, while calcium ions are known to be activatory, they can, in some cases, have a negative effect on lipase stability, as seen for *Candida antarctica* lipase A.^35^

More recently there has been a surge of interest in the activation of enzymes in neoteric solvents such as DESs.^14,36–38^ A DES is a eutectic mixture, typically formed through interactions between hydrogen bond donors (HBDs) and hydrogen bond acceptors (HBAs). Common HBAs are ammonium or phosphonium-based salts and common HBDs are urea, glycerol, ethylene glycol, and glucose.^36,38^ Perhaps the most widely used DES is that composed of choline chloride and glycerol^5^ typically used at a mole ratio of 1:2 (choline:glycerol). This DES is widely used due to its relatively low viscosity.^39^

In addition to being cheap, tunable, easily synthesised and environmentally benign, perhaps the most interesting aspect of DES is their ability to stabilise and support enzyme catalysis without loss of enzyme activity. This makes DESs extremely attractive as solvents for biocatalytic conversions. Indeed, the most extraordinary aspect of biocatalysis in DESs is the common observation that, unlike organic solvents, enzymes are not only soluble in DES but, in a surprising number of cases, they are activated and stabilised (see Table 1). This ability of DES to activate and stabilise enzymes is unique in biological chemistry and offers the potential to unlock significant reaction efficiency with the resulting reduction of costs. This activation is mainly observed in DES based on cholinium salts as hydrogen bond acceptors (Table 1). The vast bulk of DESs employed to date have been polar solvents although the use of hydrophobic DES (e.g. lidocaine-based hydrophobic DES) is increasing and has been successful for some biocatalytic applications.^40^ Thus, DESs appear to be a universal “key” to unlock the catalytic potential of several enzymes used in biocatalysis. This suggests some commonality of mechanism that is incompletely understood at present. It should be noted that the use of DES media to support biocatalysis entails exposing enzymes to solutes (choline chloride, glycerol, glucose, *etc.*) at levels that are much higher than these enzymes would encounter in their natural state in the cell. For example, choline is found intracellularly at levels of 15 µM but in DES, levels can be in the mM range and higher depending on the ratio of DES to aqueous phase. It might be assumed that the altered properties of enzymes in DES (activation, stabilisation) are a function of their exposure to such high solute concentrations. In previous studies, lipase activation was explained as being due to either, a specific charge interaction^41^, hydrogen bonding to activate the enzyme by promoting a more active conformation or a direct interaction with the acyl-binding pocket.^42^

**Table 1.**
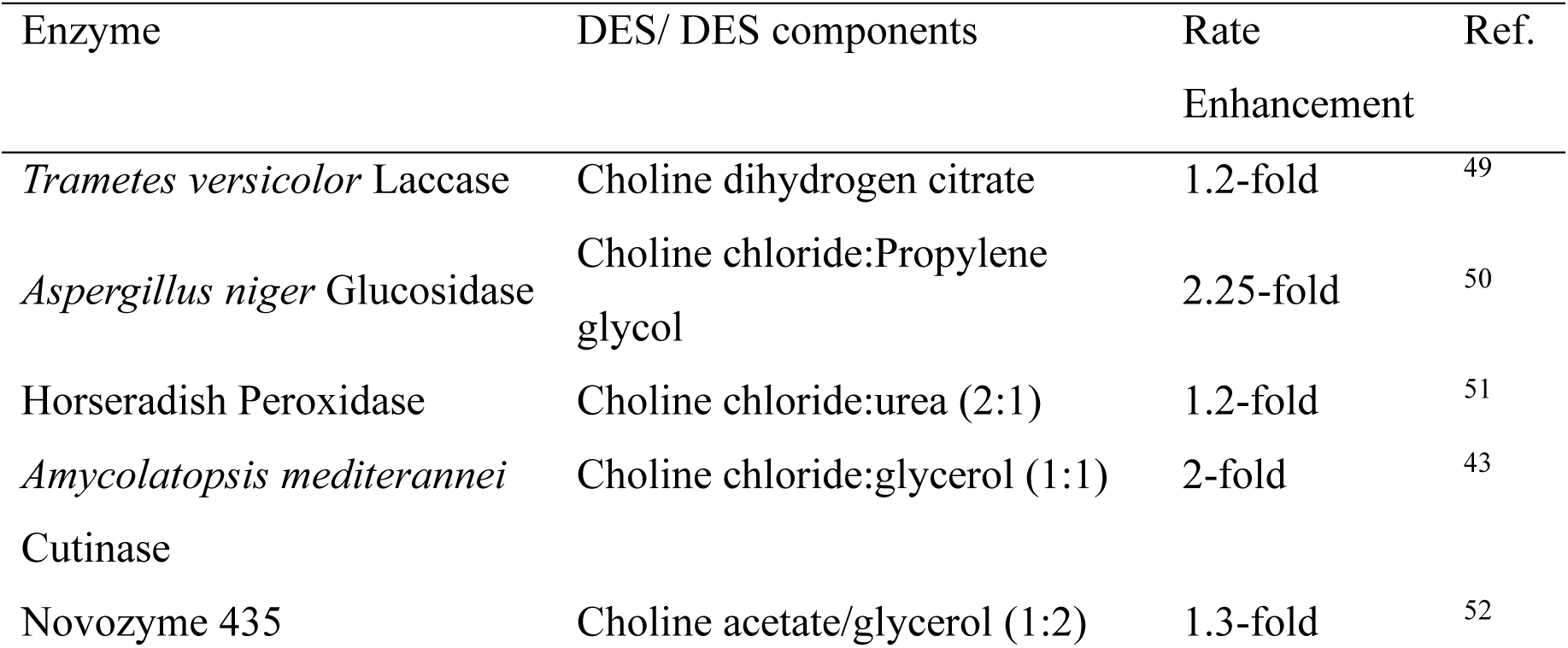
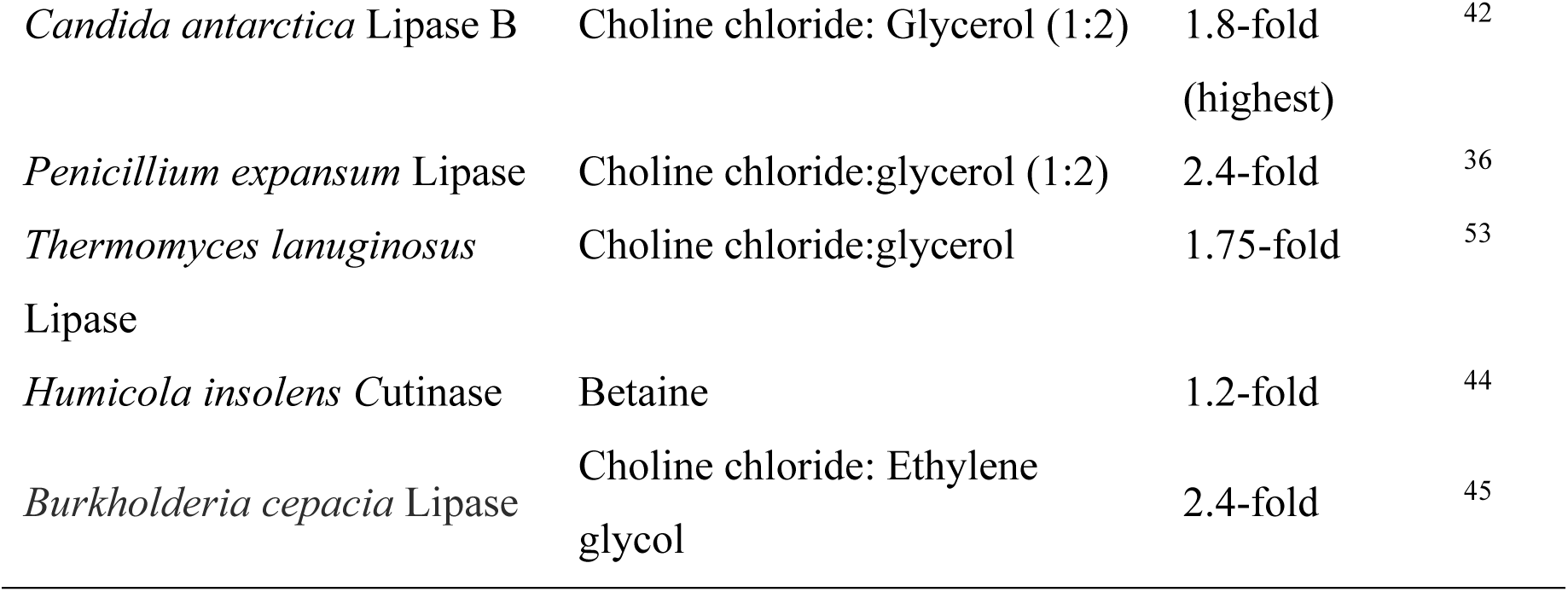
Enzymes used for biocatalysis that are activated in Deep Eutectic Solvents. Note: in all cases, activity increases were accompanied by stability enhancement.

| Enzyme | DES/ DES components | Rate Enhancement | Ref. |
| --- | --- | --- | --- |
| *Trametes versicolor* Laccase | Choline dihydrogen citrate | 1.2-fold | 49 |
| *Aspergillus niger* Glucosidase | Choline chloride:Propylene glycol | 2.25-fold | 50 |
| Horseradish Peroxidase | Choline chloride:urea (2:1) | 1.2-fold | 51 |
| *Amycolatopsis mediterranei* Cutinase | Choline chloride:glycerol (1:1) | 2-fold | 43 |
| Novozyme 435 | Choline acetate/glycerol (1:2) | 1.3-fold | 52 |
| *Candida antarctica* Lipase B | Choline chloride: Glycerol (1:2) | 1.8-fold<br>(highest) | 42 |
| *Penicillium expansum* Lipase | Choline chloride:glycerol (1:2) | 2.4-fold | 36 |
| *Thermomyces lanuginosus*<br>Lipase | Choline chloride:glycerol | 1.75-fold | 53 |
| *Humicola insolens* Cutinase | Betaine | 1.2-fold | 44 |
| *Burkholderia cepacia* Lipase | Choline chloride: Ethylene glycol | 2.4-fold | 45 |

The effect of DES on enzymes has been described as a “supramolecular net”.^38,41^ As we and others have pointed out, however, many reports of activation in DES are in fact, activation in dilute DES component solutions.^9,43–45^ The dilution is often so high that the nanostructure of the DES is dispersed. Thus, as we have argued in previous work,^41^ the enhancements observed in DES do not consider its dissociation when dispersed in buffer solutions. In brief, DES retains its nanostructure properties after mixing with aqueous solutions to an extent that depends on the ratio of water to DES. At high water levels, a DES solution simply becomes an aqueous mixture of its components. There are essentially three phases when water or aqueous buffer is mixed with DES. At low levels of water (0-30% w/w), the added water is dispersed within the DES network of hydrogen bonds. At this level, the DES retains its essential solvent properties. At higher levels of water (30-50% w/w), the DES network is disturbed, and it is fragmented into clusters within an aqueous phase. At even higher water levels (above 50% w/w), DES components are simply dissolved solutes in an aqueous medium.^46,47^ At the highest water levels, where a DES does not retain its nanostructure, the effect of DES on biocatalytic reactions should be primarily due to its individual components rather than a bulk solvent effect.

Strikingly, several enzymes, including lipases and closely related cutinases are activated in DES solutions (see Table 1). CALB, for example, is activated by choline based DES media.^42,48^ It is, perhaps, somewhat surprising that lipases, usually known for activation via hydrophobic interactions, can be activated in polar DES solvents. A central unanswered question concerning the activation of lipases in DES is to reconcile the various avenues for activation. How is it possible that a lipase may be activated by interfacial activation, by attachment to hydrophobic supports, by detergents, by calcium salts, and by polar eutectic solvents? How are these modes of activation related to one another, and can they be exploited in concert to achieve even greater catalytic potential? The first, second and third imply a hydrophobic interaction while the last two imply a possible hydrogen bonding or charge interaction. There is a need for clarity to advance our understanding of this area since these interactions have the potential to unlock the latent catalytic potency of a biocatalysts.

Previous work in this laboratory has revealed the central role of choline in the activation of alcohol dehydrogenase and cutinase enzymes. Herein, we show that choline is, indeed, an activator of CALB and show that the activation is due to the quaternary ammonium moiety of choline. This allowed us to discount the hydrogen bonding theory or the formation of a supramolecular net as activation mechanisms. Furthermore, we show that the hydrophobic alkyl corona surrounding the nitrogen ion greatly influences the degree of activation. Surprisingly, increasing the hydrophobicity around the quaternary nitrogen moiety by increasing alkyl chain length can provide extraordinary activation of CALB. For instance, tetraoctylammonium bromide showed an ∼ 5-fold activation of CALB at low micromolar (µM) levels (soluble form). This is, to the best of our knowledge, the highest degree of activation for a small molecule, non-specific activator, reported to date, for this or any enzyme.

Herein, we also demonstrate that CALB, like many lipases, is activated by calcium. Mixed activator experiments using calcium ions and choline showed the activation of CALB can proceed through binding at two separate and independent sites on the enzyme. These studies raise some important questions regarding the lid-opening model as the sole route to lipase activation. The different sites for activation either both act to open the “lid” or there is more than one pathway to lipase activation. The fact that we had two soluble activators apparently acting independently was intriguing. Activation by calcium salts destabilised CALB while ChCl activation was stabilising – thus, activation at one site (alkylammonium binding site) provides enhanced stability over binding at another (calcium binding site).

This work establishes these alkylammonium salts as a new class of activator for biocatalysis that are applicable to a wide number of enzymes. Importantly, the mechanism of activation described herein provides an explanation that reconciles the various types of CALB activation. Thus, it shows that the activation seen in DES, the activation upon immobilisation on hydrophobic supports and the activation by detergents with long alkyl chains can all be traced to a hydrophobic patch on the enzyme surface. Activation occurs when this site is in contact with a hydrophobic entity, whether it is an immobilised support^54,55^, a component of DES^42^, or a detergent with a cationic polar head group.^28^

Alkylammonium ions, as a new class of enzyme activator, have the important benefit of being water soluble compounds that can be used as a basis for optimisation of biocatalytic activity enhancement and to probe the mechanism of activation. The data herein, along with that of other studies, allows us to explain the nature of the activation by DES and to suggest that this is a widespread phenomenon that deserves greater attention for both industrial and biomedical applications.

## 2 Materials and Methods

### 2.1 Materials

Lipase enzyme from *Candida sp.* or *Candida antarctica* lipase B (CALB) – product code L3170 was procured from Merck. Choline chloride (99%) and glycerol (99+%) were purchased from Fisher Scientific. Paranitrophenyl acetate (pNPA), Tris-base (99+%), tetramethylammonium bromide (TMAB), tetraethylammonium bromide (TEAB), tetrabutylammonium bromide (TBAB), tetrabutylammonium chloride (TBAC), tetrahexylammonium bromide (THAB), tetraoctylammonium bromide (TOAB), monomethyltricotylammonium chloride (MTOAC), tetrakis(decyl)ammonium bromide (TDAB), were obtained from Merck. All the chemicals were of analytical grade and used as received without further purification. Ultrapure de-ionised water (< 15 MΩ.cm) was used to prepare all the solutions.

### 2.2 Methods

#### 2.2.1 Lipase activity

The activity of lipase was measured by monitoring the hydrolysis of paranitrophenyl acetate (pNPA) with the liberation of paranitrophenol (pNP). A stock solution of 100mM pNPA was prepared in acetonitrile. A 1mg/mL enzyme solution was prepared in 50 mM Tris-HCl buffer, pH 7.8. For the assay, solution A was prepared by mixing 50 mM Tris-HCl buffer with ethanol and pNPA substrate at a ratio of 95:4:1. The reaction was started by adding 20 µL of enzyme solution (1mg/mL) to 230 µL of solution A in a 96-well plate (Greiner 96well multiplate with solid U bottom - 650161G), such that the final assay mixture contains 1mM pNPA, 1% (v/v) acetonitrile and 4% (v/v) ethanol in 50 mM Tris-HCl buffer. The assay mixture was incubated at 35°C and the absorbance at 400 nm was measured at 5.0 min intervals using a Thermo Scientific 215 Varioskan® LUX microplate reader. Product formation was followed by monitoring the formation of pNP (400 nm) and pNP concentration was estimated by using a standard calibration plot. Throughout this report, the pNP concentration formed after 20 mins of hydrolytic reaction was used. Owing to the autohydrolysis of substrate (pNPA), a substrate blank (containing substrate without enzyme in buffer with salt) was performed for each experiment and its absorbance was subtracted from the sample absorbance.

To test the effect of various salts in the hydrolytic activity of CALB, different concentrations of ChCl, Gly, and tetraalkylammonium salts were prepared in 50mM Tris-HCl buffer (pH 7.8) and used in the assay. Stock solutions of 3.0 M of ChCl, 1.5 M of TMAB and TEAB, 1.0 M of TBAB and TBAC were prepared. Due to solubility limitations for the higher alkyl chain salts, stock solutions of 10.0 mM for THAB, 150 µM for TOAB and TDAB. For MTOAC salt, stock solution of 3.0 mM was prepared. Note that all these salt solutions were freshly prepared and used within a day. The pH of all solutions was maintained at 7.8 using 50mM Tris-HCl buffer, unless specified otherwise. The activity of CALB for each reaction was expressed as a percentage, relative to a control (containing substrate and enzyme in the buffer solution, without TAA salt or effector).

#### 2.2.2 Statistical analysis

Triplicates of each sample were performed in a single measurement; and each experiment was performed a minimum of three times (n=9). One-Way ANOVA and post-hoc Tukey tests was performed on the experimental data to examine the differences among the samples using IBM SPSS Statistics 29.0.2.0.^56^ The graphs are plotted using Origin pro 2016 software. The statistical significance of difference in relative activity is indicated using the asterisk symbol above the relevant bars. Here, * represents a p value < 0.05 for a confidence interval of 95%, ** represents p value < 0.01 for confidence interval of 99 % and *** represents p value < 0.001 for confidence interval 99.9 %.

#### 2.2.3 Computational Analysis

The protein sequence of CALB was retrieved from the UniProt database (UniProt ID: P41365) and its 3D structure from the PDB (PDB: 5a71).^60^ Negatively charged, solvent accessible, CALB residues were identified using PISA^51^ and PyMol 3.0.3.^57^

*Surface Accessible Charge*: The possible binding site identification centered on solvent accessible charged residues. Molecular docking using AutoDock4^58^ was performed between different alkylammonium ions and chain B of CALB (PDB: 5a71). The receptor and TAA were pre-processed using MGLs’ AutoDock tools to add polar hydrogens, Kollman charges and to assign torsion angles^59,60^. The binding site grid box was visually defined using AutoDock tools 1.5.7 with size dimensions of 40 × 40 × 40. The xyz coordinates as grid centres were set for Asp145 at 3.621, 34.67, 19.291 as in Figure 7 and Figure 8.

Discovery Studio v24.1.0 was used to visualize possible 2D interactions and PyMol 3.0.3 for 3D poses between the ligand and the enzyme.^61^

## 3 Results and Discussion

The components of the most widely used DES are choline chloride (ChCl) and glycerol (Gly) at a mole ratio of 1:2.^42^ Based on our previous observations, the activation of enzyme by DES was traced to the choline component of DES.^41,43^ In the present work, the effect of Choline Chloride (ChCl) and Glycerol (Gly) on CALB activity was investigated in the first place. Subsequently, the effect of chemical species with structural similarities to ChCl were explored. The effect of divalent metal ions on CALB activation was also examined.

### 3.1 Effect of DES components on CALB

Figure 1 (a) shows the % relative activity of CALB in the presence of choline chloride (ChCl) over a concentration range from 5.0 mM to 3.0M. To accommodate the extended concentration range, the x-axis is logarithmically scaled, and the control (100%) is indicated as a dotted horizontal line.

**Figure 1:**
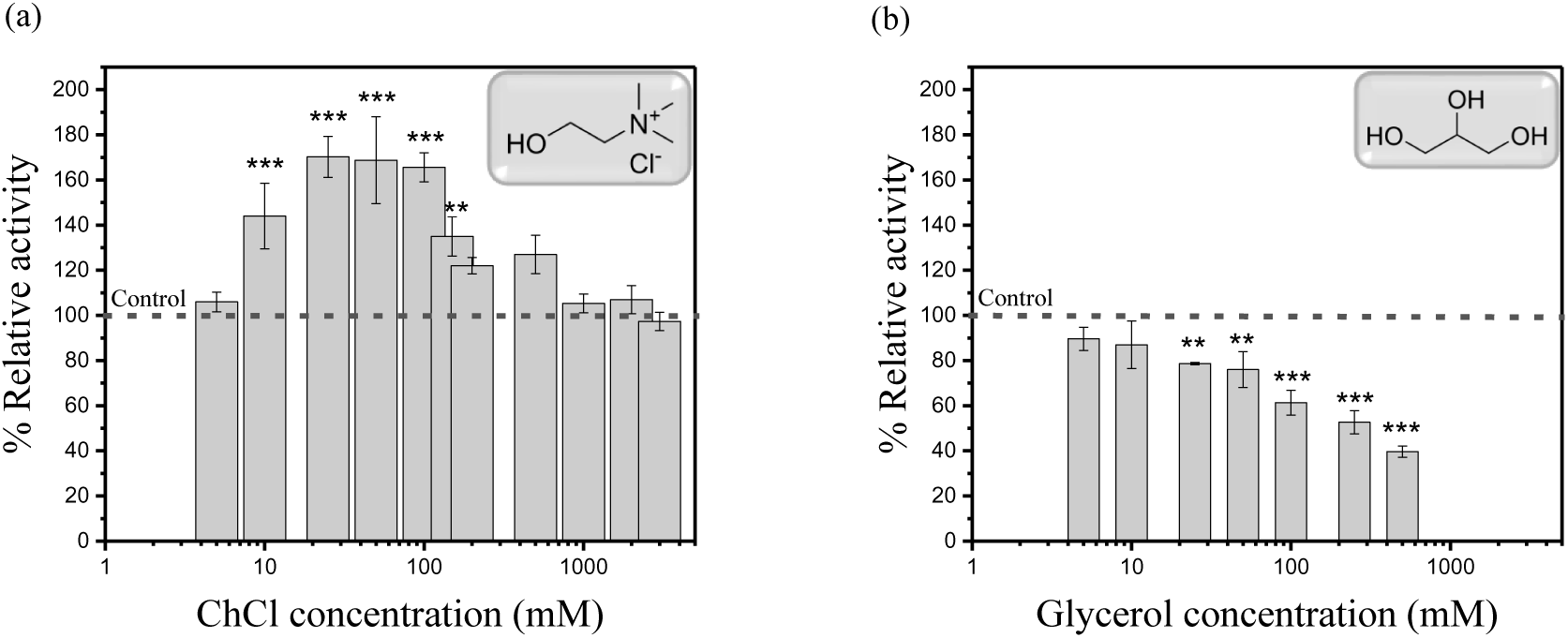
(a) shows the effect of ChCl on CALB activity from 5.0mM to 3.0M and (b) shows the effect of Glycerol on CALB activity over the concentration range from 5 to 500mM. The x-axis is a logarithmic scale, the control (100%) is indicated as a dotted horizontal line and the inset shows the chemical structure of the corresponding test molecule. The assay used was the standard assay (see methods) and the substrate was pNPA at a concentration of 1.0mM. All assays were temperature-controlled at 35°C. ** represents p value < 0.01 for confidence interval of 99 % and *** represents p value < 0.001 for confidence interval 99.9 %.

The data of Figure 1 (a) clearly shows a 1.7-fold activation (170%) of CALB by choline. This is slightly higher than the activation of CALB (157%) reported in a DES composed of choline chloride:glycerol.^62^ This finding proves that CALB can be activated by ChCl alone to the same extent as observed in a DES solution. Figure 1 (b) shows that glycerol inhibited CALB even at the lowest concentration used here (5.0 mM).

These findings add to the growing body of evidence that points to the choline component of DES as the enzyme-activating agent in DES media. The activity of CALB in DES is, therefore, the resultant of its activation by ChCl and its inhibition by Gly. This is the third example of ChCl activation we have observed in this laboratory.^41,43^ This is the first demonstration of this type of activation for CALB and is the first report of CALB activation by an intracellular compound.

It was notable that above 100mM ChCl, CALB activity began to decline (see Figure 1 (a)). When ChCl levels in a DES are too high, inhibition is observed. This profile of activation followed by inhibition is like that observed for Horse Liver Alcohol Dehydrogenase.^41^ Inhibition of CALB by higher levels of ChCl was reported previously.^62^ It is often reported that DES-mediated activation of enzymes occurs in a mixture of DES and buffer wherein the DES is added as a minor component, as a co-solvent. At higher levels, therefore, DES is not effective as an activator but becomes an inhibitor.

We examined CALB activation by structurally related HBAs. The structures of the salts used in this study are shown in Figure 2.

**Figure 2:**
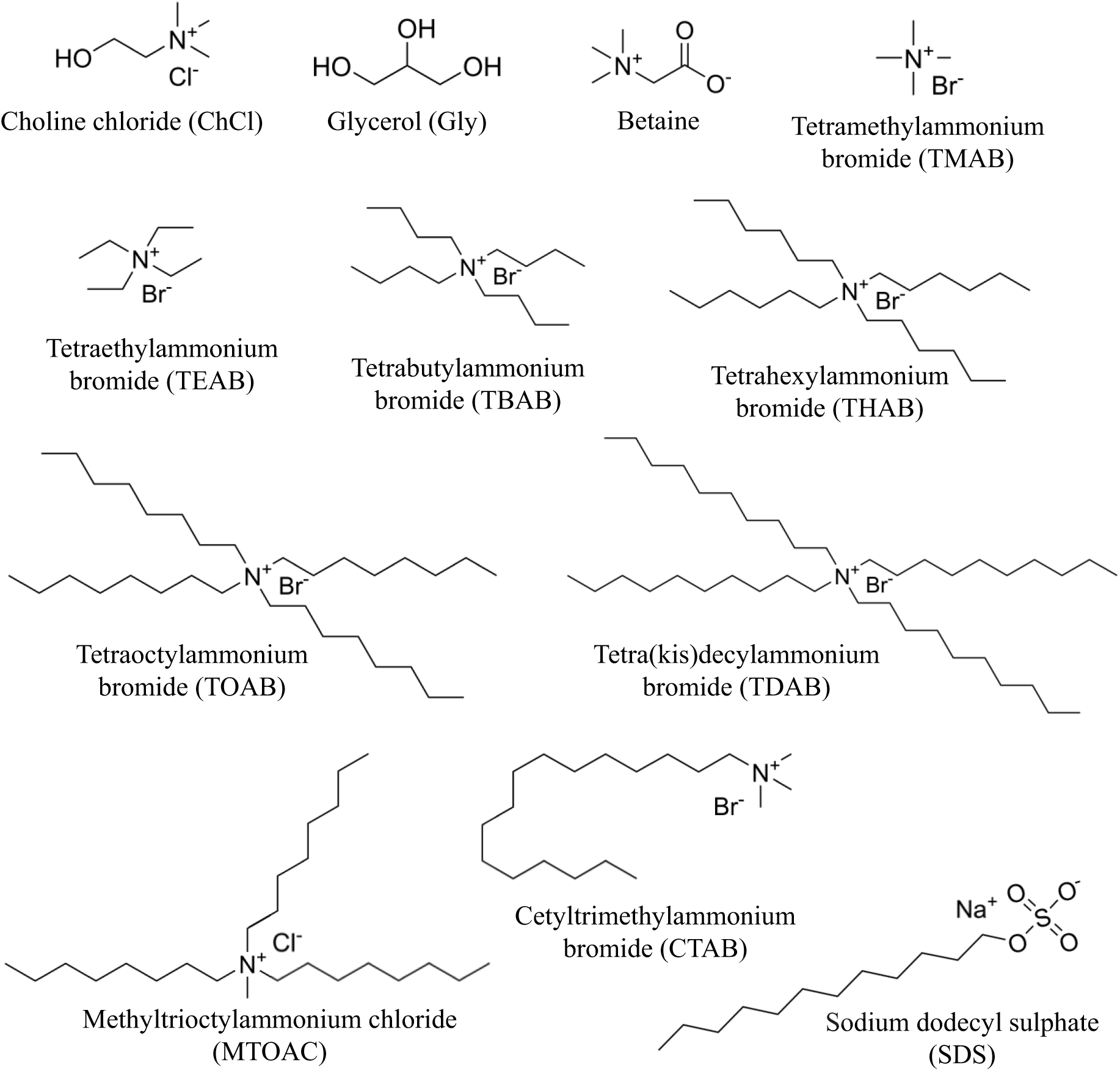
Structures of various compounds used in this study to investigate their effect on CALB activity

ChCl is a trimethylammonium (-N^+^(CH_3_)_3_) salt with a hydroxyethyl (-CH_2_-CH_2_-OH) substituent. To examine the influence of these two functional groups, we investigated the effect of the trimethylammonium ion and betaine (trimethylglycine) on CALB activity. The effect on CALB activity is shown in Figure 3. For TMAB, (Figure 3 (a)), activation is observed for concentrations higher than 500mM, but the activation is not as high as that observed with ChCl at 50mM. Note, sodium chloride (NaCl) at concentrations up to 100mM had no significant activating effect on CalB (supplementary information Figure S1). Clearly, the presence of the hydroxyl group on choline has a role in activation. By contrast, betaine inhibited CALB over a similar concentration range (Figure 3 (b)).

**Figure 3:**
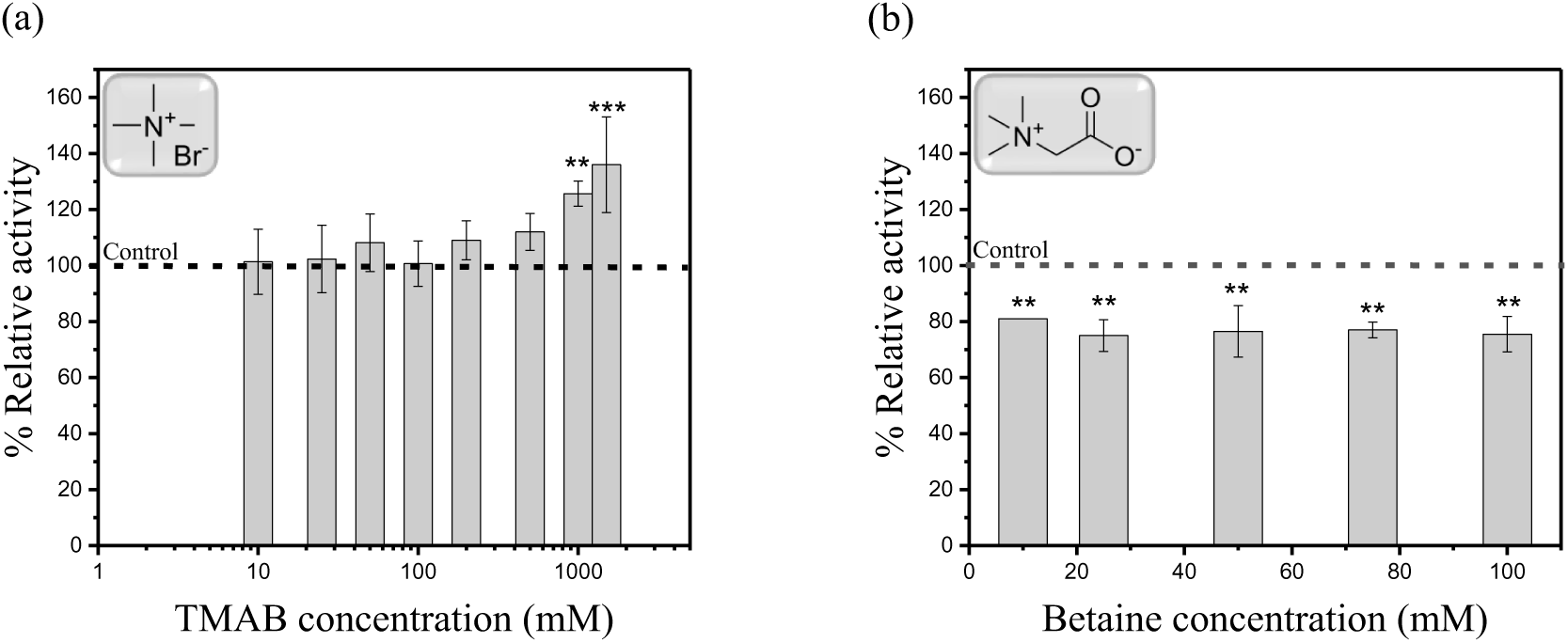
(a) shows the effect of TMAB on CALB activity from 10mM to 1.5M (x-axis is a logarithmic scale) (b) shows the effect of betaine on CALB activity from 10 to 100mM (x-axis is a linear scale). The control (100%) is indicated as a dotted horizontal line and the inset shows the chemical structure of the corresponding salt. The assay used was the standard assay (see methods) and the substrate was pNPA at a concentration of 1.0mM. All assays were temperature controlled at 35°C. ** represents p value < 0.01 for confidence interval of 99 % and *** represents p value < 0.001 for confidence interval 99.9 %.

Having established that TMAB activates CALB, we examined a series of medium and long chain structural analogues of this compound to explore the relationship between structure and degree of activation.

### 3.2 Effect of quaternary alkylammonium salts on CALB

Tetraalkylammonium (TAA) salts with increasing alkyl chain length, on the quaternary nitrogen, were examined. Figure 4 shows the effect of the medium chain TAA salts, TEAB and TBAC on CALB activity.

**Figure 4:**
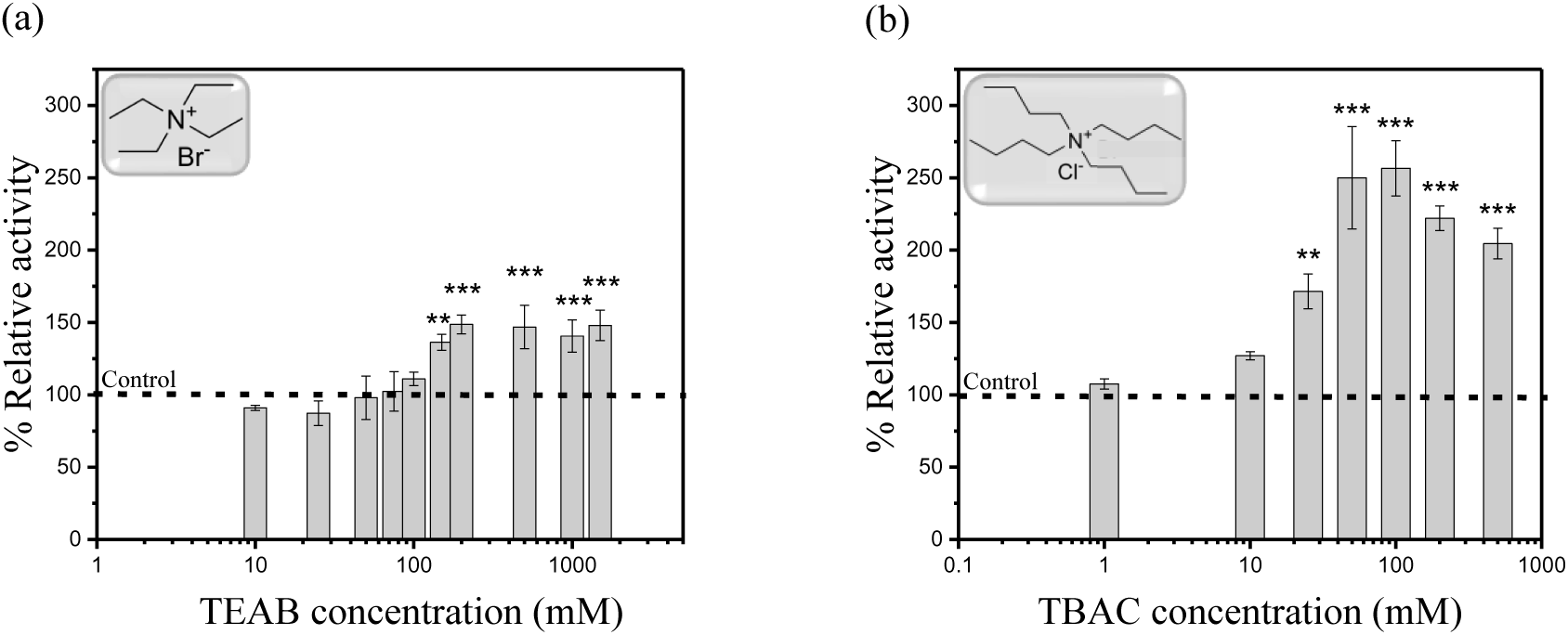
Profile of CALB activity in the presence of varying amounts of alkylammonium salts. (a) TEAB (tetraethylammonium bromide), x-axis in logarithmic scale and (b) TBAC (tetrabutylammonium chloride), x-axis in logarithmic scale. The control (100%) is indicated as a dotted horizontal line and the inset shows the chemical structure of the corresponding salt. The assay used was the standard assay (see methods) and the substrate was pNPA at a concentration of 1.0mM. All assays were temperature-controlled at 35°C. * represents the p value < 0.05 for a confidence interval of 95%, ** represents p value < 0.01 for confidence interval of 99 % and *** represents p value < 0.001 for confidence interval 99.9 %.

Figure 3 (a) showed that TMAB alone was sufficient to activate CALB albeit to a lesser extent than choline. Both the medium chain alkylammonium salts activated CALB as seen in Figure 4 (a) and (b) and the trend showed increasing activation as alkyl chain length increased. A maximum of 150 % activation was observed for TEAB at concentrations > 250 mM whereas, CALB showed a 2.5-fold (250%) activation in the presence of 50mM TBAC. To consider the effect of the anionic counterion, the chloride salt of tetrabutylammonium (TBAC) was tested and its effect on CALB is shown in Figure 4 (b). The effect by its corresponding bromide salt (TBAB) is shown in supporting information Figure S2. The % activation achieved by the bromide salt is the same as for the chloride salt.

Based on these observations, it is clear that an enhanced hydrophobic interaction is the most significant factor in CALB activation. This profile is like that observed for horse liver ADH.^52^ For significant activation of CALB, the required concentrations of medium chain length TAA salts is in the millimolar (mM) region. Further investigation explored the effect of increasing alkyl chain length on CALB activity. As the chain length of TAA salts increases, they become less soluble in aqueous buffer and millimolar solutions are not possible to prepare. Stock solution of these salts were prepared as detailed in section 2.2.1.

Figure 5 shows the effect of long chain TAA salts (tetrahexyl to tetradecyl) on CALB activity. It is noteworthy that these compounds activate CALB at concentrations in the micromolar (µM) region. THAB showed 2.3-fold activation (230 %) of CALB at 50 - 100 µM concentration. TOAB, is particularly potent, showing activation of ∼5-fold (512 %) at 100 µM concentration. This derivative was relatively tight binding and a Kd of ∼5×10^−6^ M was estimated from the data of Figure 5 (b). Further increasing the alkyl chain length (decyl, 10 carbons) showed only a ∼2 fold increase (196 %) in CALB activity at 100 µM. The poorly soluble tetradecyl salt (TDAB) does activate, but not to the same extent as the TOAB salt. The tetradecyl moiety may sterically exclude its binding, thereby, providing lower activation than TOAB.

**Figure 5:**
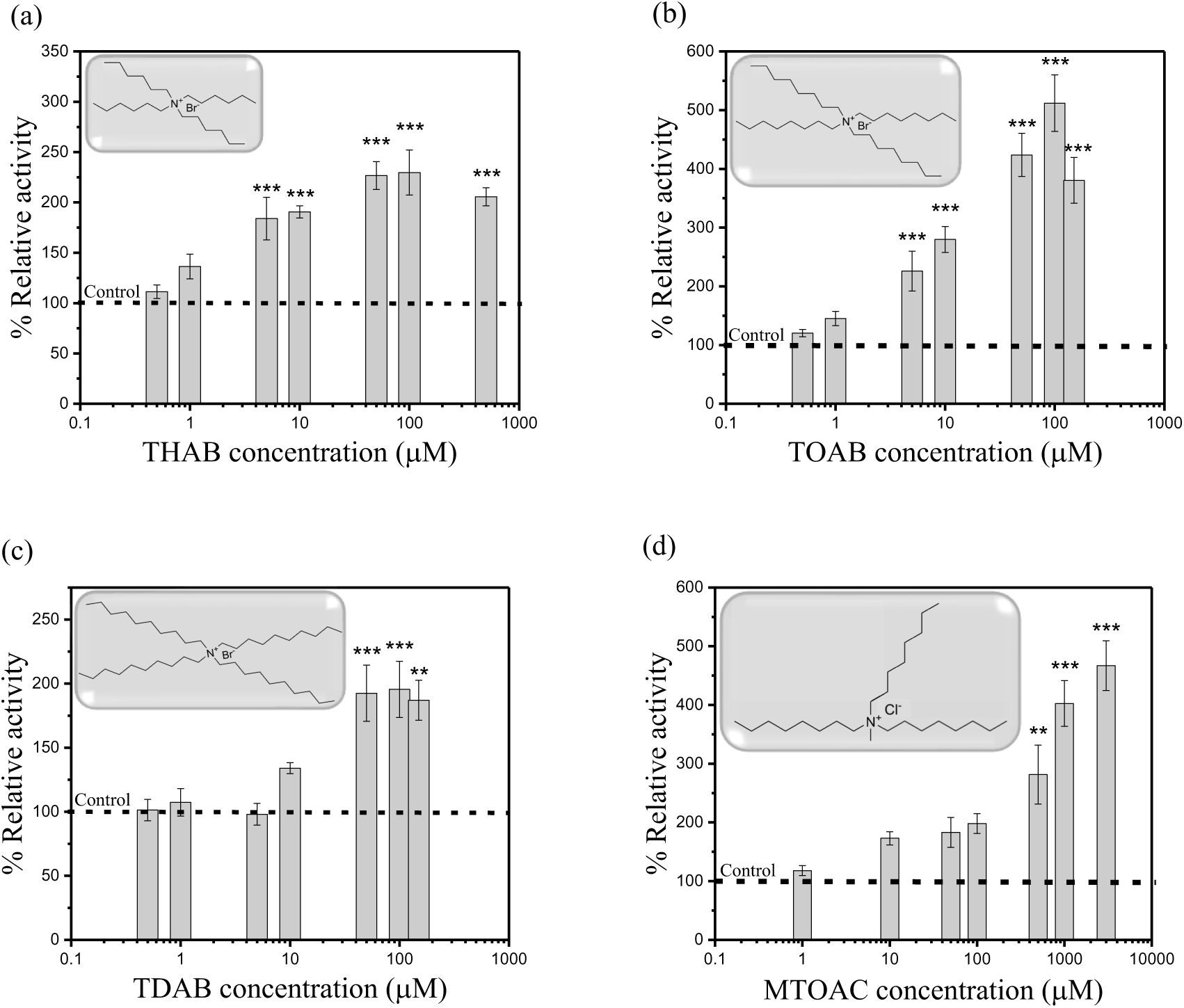
Cal B activation by long chain alkylammonium salts. (a) THAB (tetrahexylammonium bromide), (b) TOAB (tetraoctylammonium bromide), (c) TDAB (tetrakis(decyl)ammonium bromide) and (d) MTOAC (methyltrioctylammonium chloride). All the plots have the x-axis as a logarithmic scale. The control (100%) is indicated as a dotted horizontal line and the inset shows the chemical structure of the corresponding salt. The assay used was the standard assay (see methods) and the substrate was pNPA at a concentration of 1mM. Note that the concentration for the observed activation moves into the micromolar range for the longer-chain salts. All assays were temperature controlled at 35°C. * represents the p value < 0.05 for a confidence interval of 95%, ** represents p value < 0.01 for confidence interval of 99 % and *** represents p value < 0.001 for confidence interval 99.9 %.

These findings seem to suggest that the alkylammonium’s positive charge serves to direct this interaction to a “landing site” on the enzyme surface. To examine the effect of the octyl group, we investigated the activation of CALB by the methyltrioctylammonium ion. The lack of this single octyl group rendered MTOAC more soluble (3.0mM) than TOAB. The profile is shown in Figure 5 (d). It is evident that, MTOAC increased CALB activity and a maximum of ∼ 5-fold activation was achieved 3.0mM.

Figure 6 is a plot of maximum activity for TAA salts as a function of alkyl substituent. This clearly shows dramatic activation by the tetraoctyl salt. Indeed, 1.0 µM TOAB had the same activating effect as 100mM choline (maximum activation level), a roughly 1.5-fold activation (Figure 1(a) and Figure 5(b)). The activation by TOAB (∼5-fold) is, to the best of our knowledge, the highest activation reported to date for a non-peptide small molecule activator. The extraordinary aspect of this finding is that this activation occurs at micromolar concentrations of the TOAB salt. The longer chain decyl salt was still a potent activator but maximum activation only reached 200%. When the methyltrioctylammonium salt (MTOAC) was tested, it showed activation of 200 % for 100 µM. This is lower than that achieved for 100 µM of TOAB salt. However, by increasing the MTOAC concentration to 3.0 mM (which is possible due to its increased solubility), the activity increased up to 470 %. This analogue, therefore, can reach high levels of activation but at much higher concentrations than required for the tetraoctyl salt. Further increasing its concentration was limited by its aqueous solubility. These observations imply that all four 8-carbon chains are needed for maximum activation at micromolar levels. Clearly, the multi-directional geometry of the tetrahedral nitrogen is important for binding.

**Figure 6:**
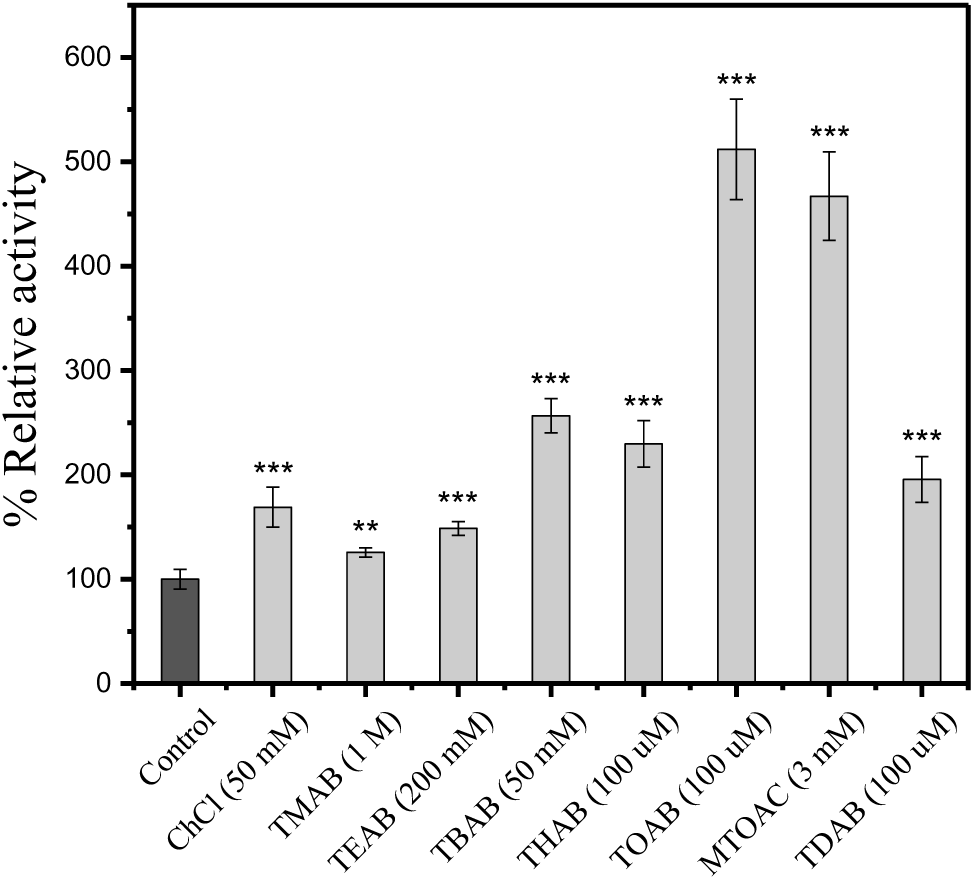
Summary of the effect of alkylammonium salts with varying alkyl substituents on the activation of CALB. The first dark gray bar indicates the control. The concentration at which maximum activation achieved for each salt is indicated on the x-axis labels within brackets. ChCl is choline chloride, TMAB is tetramethylammonium bromide, TEAB is tetraethylammonium bromide and TBAB is tetrabutylammonium bromide, THAB is tetrahexylammonium bromide, TOAB is tetraoctylammonium bromide, MTOAC is methyltricotylammonium chloride and TDAB is tetrakis(decyl)ammonium bromide. * represents the p value < 0.05 for a confidence interval of 95%, ** represents p value < 0.01 for confidence interval of 99 % and *** represents p value < 0.001 for confidence interval 99.9 %.

These observations allow us to make certain deductions. The presence of the charge on alkylammonium ions is not the main determinant of activation, it is clearly the hydrophobic corona surrounding the quaternary nitrogen that influences the degree of activation. However, the charge serves to orient the hydrophobic component towards the binding site. TAA salts are well known in synthetic organic chemistry as phase transfer catalysts. It might be assumed that their action is due to an effect on solvent parameters, or due to hydrogen bonding with the enzyme surface or even a detergent effect.^62,63^ However, the fact that the longer chain derivatives activate at such low concentrations allows us to discount these explanations. Activation cannot involve micelle formation for these salts since they do not form micelles at such low concentrations. Two recent studies have observed activation of CALB in DES.^42,62^ Those studies explained the activation of CALB on the basis of hydrogen bonding interactions and/or charge interactions with surface residues on CALB.

If we consider the activation seen for other enzymes using these salts, we can conclude that this type of activation is widespread and potentially can be exploited for a wide variety of benefits. A surprising number of enzymes are reported to be activated in DES solutions. These include examples of proteases, lipases, cutinases, dehydrogenases, peroxidase etc. (see Table 1). In almost all cases, activation was seen mostly for the cholinium salts. To date, we have evidence that several of these could actually be activated by choline alone. Herein, we suggest that the mechanism of activation is likely to be the same for many of these enzymes, namely, binding of the cholinium ion to a specific region on the enzyme surface. This reveals this class of compounds have a broad utility as enzyme activators across a range of enzyme classes.

### 3.3 Molecular docking

#### 3.3.1 Exploring a possible alkylammonium binding site on CALB

Molecular docking can provide valuable insights into the interaction between the activating compounds and CALB, revealing details such as predicted binding site interactions, and binding energy.^59^ The structure of the TAAs carries a permanent positive charge so it likely interacts with a negative charge on the surface of CALB. Possible interactions can be due to a direct interaction between the quaternary ammonium group and a negatively charged amino acid or between the aliphatic chains and adjacent hydrophobic amino acids.

Due to the small size of these TAA compounds several potential binding sites might be expected given that the surface of the enzyme has 17 (see supplementary information TableS1) surface exposed negative charges (aspartate and glutamate) on chain B of CALB (pdb:5a71). Chain B is chosen, since this is reported as a closed monomer of CALB based on X-ray crystal structures^24^. Identifying a precise binding location for the activating compounds is challenging. Figure 7 shows the molecular docking of different TAA ions with the surface of CALB. The pose with the lowest predicted binding energy (greater affinity) involved an attractive charge interaction with Aspartate-145 (Asp145) and predicted binding scores are shown in Table 2. The computed binding energies correlate with the degree of activation observed experimentally. Asp145 is surrounded by a number of hydrophobic amino acids, which are surface accessible (see supplementary information TableS2). Hydrophobic interactions are possible with nearby residues; Val141, Leu147, Val154 and Trp155 (see surface accessible area in supplementary information TableS1).

**Figure 7:**
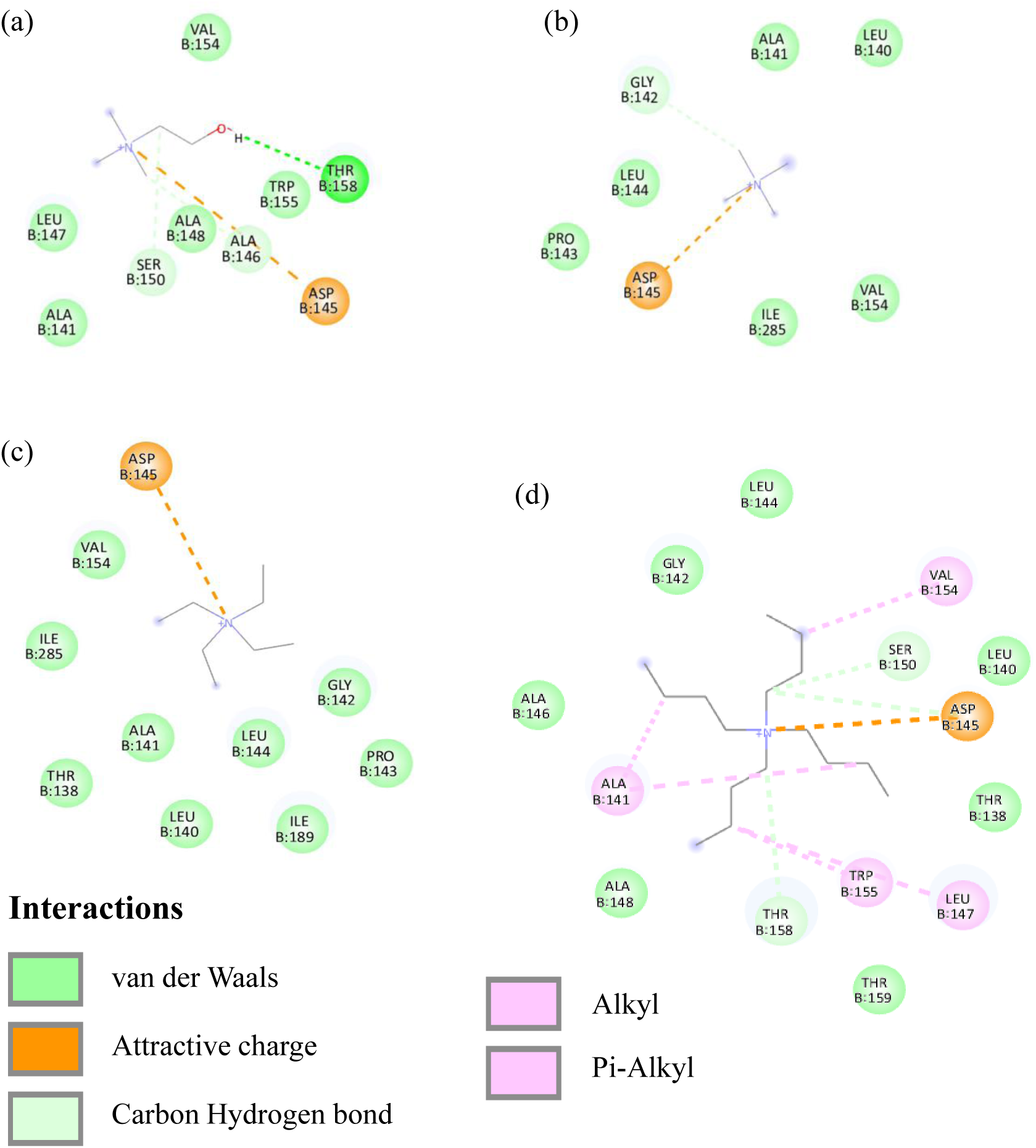
2D interaction plot between chain B (closed conformer; pdb:5a71) and different quaternary ammonium compounds a) choline, b) tetramethyl, c) tetraethyl and d) tetrabutylammonium.

**Table 2.** Binding energies for different tetraalkylammonium compounds docking with Chain B of CALB (pdb:5a71). The procedure was as outlined in materials and methods above. The estimated binding energies reflect the degree of experimentally observed activation.

| Compound | Binding energy<br>(KJ/mol) | Activation<br>(Experimental) |
| --- | --- | --- |
| Choline | -4.07 | 1.7 fold |
| Tetramethylammonium | -3.01 | 1.2 fold |
| Tetraethylammonium | -3.83 | 1.5 fold |
| Tetrabutylammonium | -5.71 | 2.5 fold |

Figure 7 shows the docking of choline, tetramethylammonium, tetraethylammonium and tetrabutylammonium ions with a surface exposed patch on CALB centered on Asp-145. Clearly, as the alkyl chain length of these TAAs increases there is greater contact with adjacent hydrophobic amino acids which enhances binding affinity. The location of this binding region (choline as ligand) relative to the active site is shown. The choline binding location (Asp-145) is roughly 14 Å from the active site.

This modelling suggests that CALB activation is likely due to docking of the quaternary nitrogen’s positive charge with Asp-145 and it is enhanced when the nitrogen substituents were bulkier – greater activation was seen with the tetrabutylammonium ion than with the tetramethylammonium ion. Clearly, the aliphatic substituents enhanced binding of TAA salts by interaction with adjacent hydrophobic residues (see Figure 8).

**Figure 8:**
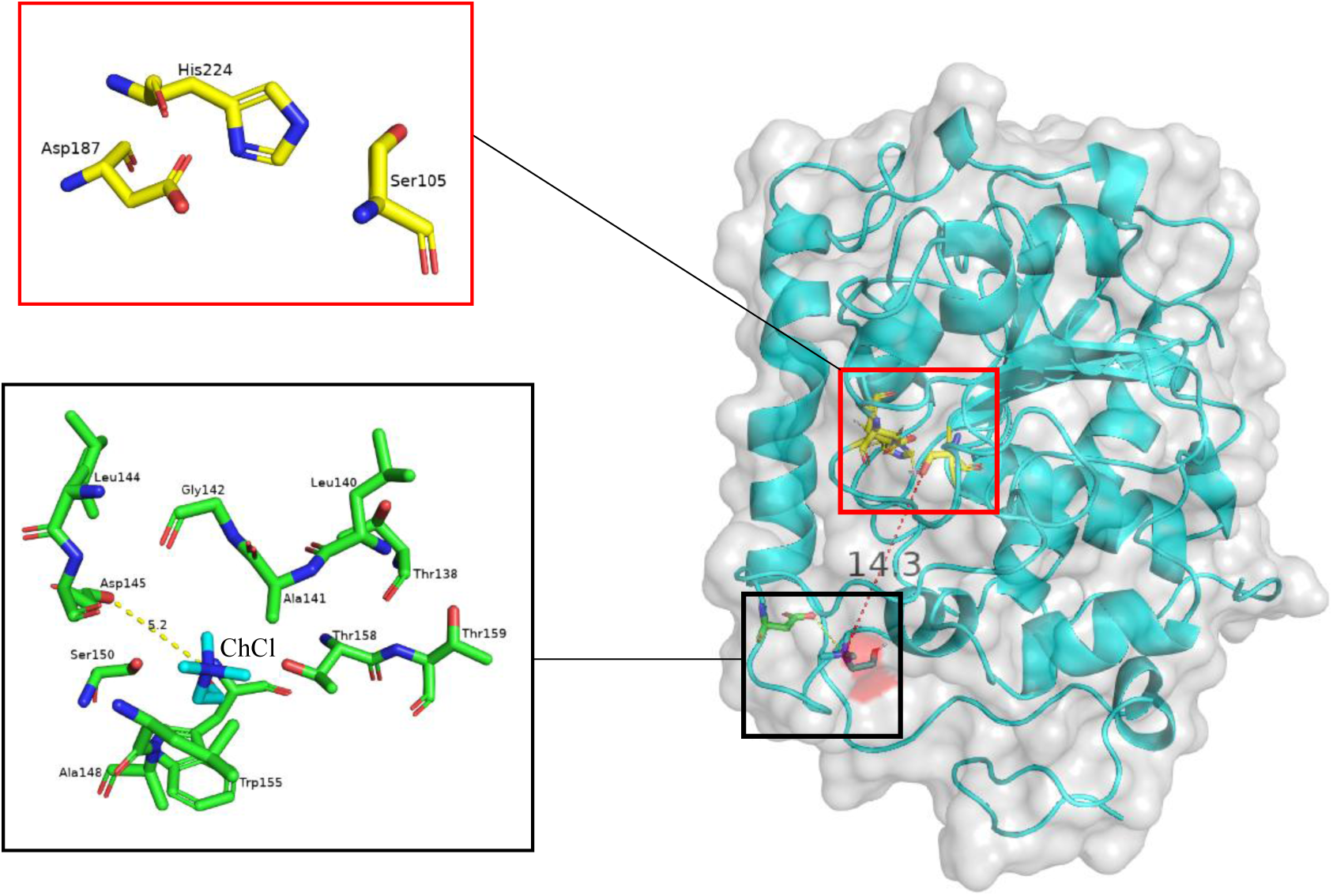
Shows the possible binding site for alkylammonium salts. Ligand (Choline) is highlighted in the black box. The distance between Asp145 and choline is 5.2 Å (yellow dotted line). The catalytic triad is shown in the red box. The distance between Asp145 and the catalytic site is approximately 14.3 Å (red dotted line).

### 3.4 Mixed activator experiment

While the above experiments explained the activation of CALB in TAA salts and identified a possible binding site, there were still unanswered questions regarding the relationship between TAA activation and activation by calcium. In this work, we investigated various divalent metal ions (Ca^2+^, Ba^2+^, Mg^2+^, and Zn^2+^) for their effect on CALB activity and the findings are shown in Supplementary information Figure S3. Among these, Ca, Ba and Mg activate CALB at a concentration of 10mM. Calcium activates CALB, reaching an activation of around 200% at 50mM (Figure 9 (a)).

**Figure 9:**
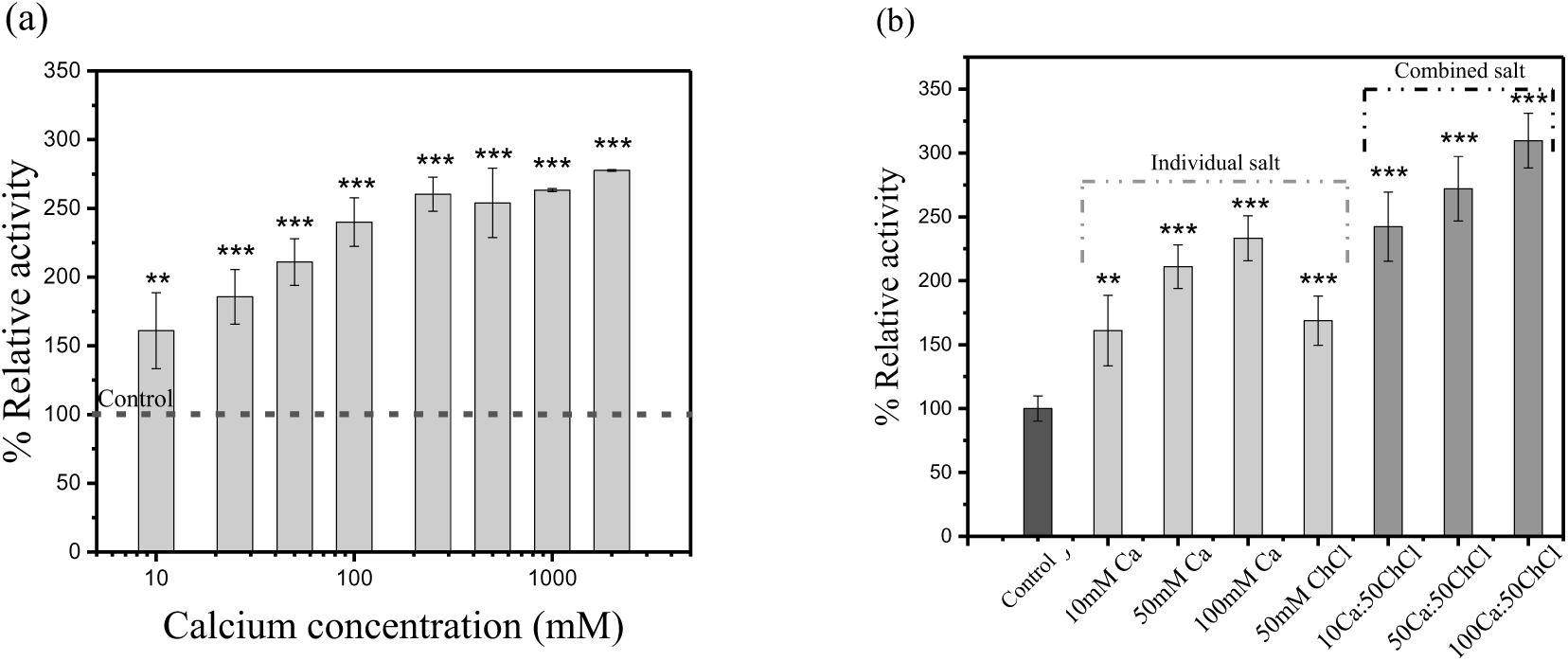
(a) Shows the activation of CALB by calcium is saturable, here the x-axis is a logarithmic scale. (b) shows the effect of combined presence of calcium chloride and choline chloride when added together to an assay mix. Note that where axis labels indicate a mixture of Calcium (Ca) and Choline (ChCl), the numerals indicate mM concentrations. * represents the p value < 0.05 for a confidence interval of 95%, ** represents p value < 0.01 for confidence interval of 99 % and *** represents p value < 0.001 for confidence interval 99.9 %.

Having two soluble activators of CALB, calcium and choline, it was of interest to compare them directly. If they were binding at the same site, they were expected to act as competitive activators. On the other hand, if these molecules could independently activate CALB, their activation might be additive. It is worth noting that activation of *Candida antarticia* Lipase A by calcium has been shown to give rise to enzyme instability.^35^ Calcium activation has been explained as calcium binding in a manner that would cause “lid” movement allowing greater access for substrate binding thereby increasing turnover.^64^

Figure 9 (a) shows a plot of CALB activation as a function of calcium concentration. This is a saturable process with maximum activity at 100mM calcium chloride. This activation was 2.3-fold over the control activity. This was somewhat higher than the 1.7-fold seen with ChCl (see Figure 1(a)). To investigate the combined effect, calcium and ChCl at varying ratios viz., 10:50, 50:50 and 100:50 (Ca:ChCl) were tested. Figure 9 (b) shows the CALB activation profile when adding calcium and choline salt together in an assay mixture. For comparison, % relative activity obtained with the individual salts in respective concentration is also shown. Surprisingly, the effect was additive where 100mM calcium chloride and 50mM choline chloride reached a combined activation of over 3-fold. Thus, the activation effect was roughly equal to the sum of each activator (calcium, choline), when used alone. On the other hand, when choline and tetrabutylammonium salt was added together, there is no significant additive effect observed, details are shown in Supplementary information Figure S4. This confirms that choline and TAA salts bind in the same location, whereas calcium and choline (and TAA) binding sites are different. This finding clearly argues for a two-site model for activation, a calcium binding site and a TAA binding site that independently activate Cal B.

### 3.5 Activation - effect on stability

In most cases in the literature, lipase activation on hydrophobic supports or in DES is accompanied by thermal stabilisation.^29^ Activation by calcium is also thought to stabilise proteins, however, in some cases, it is reported to destabilise lipases.^65^ Figure 10 compares CALB thermal stability at 50°C in the presence of ChCl and in the presence of CaCl_2_. As reported for CALA, the calcium salt caused CALB instability.^35^ Thus, the addition of calcium chloride caused CALB activity to drop to 20% of its initial value within 30 min at 50 °C. The enzyme control lost ∼50% of its activity under the same conditions. The presence of ChCl dramatically improved thermal stability of CALB and lost only ∼ 20% activity in 30min. This finding further argues for a two-site mechanism for CALB activation. The activation via hydrophobic interaction has greater stabilising effects than that at the metal ions binding site.

**Figure 10:**
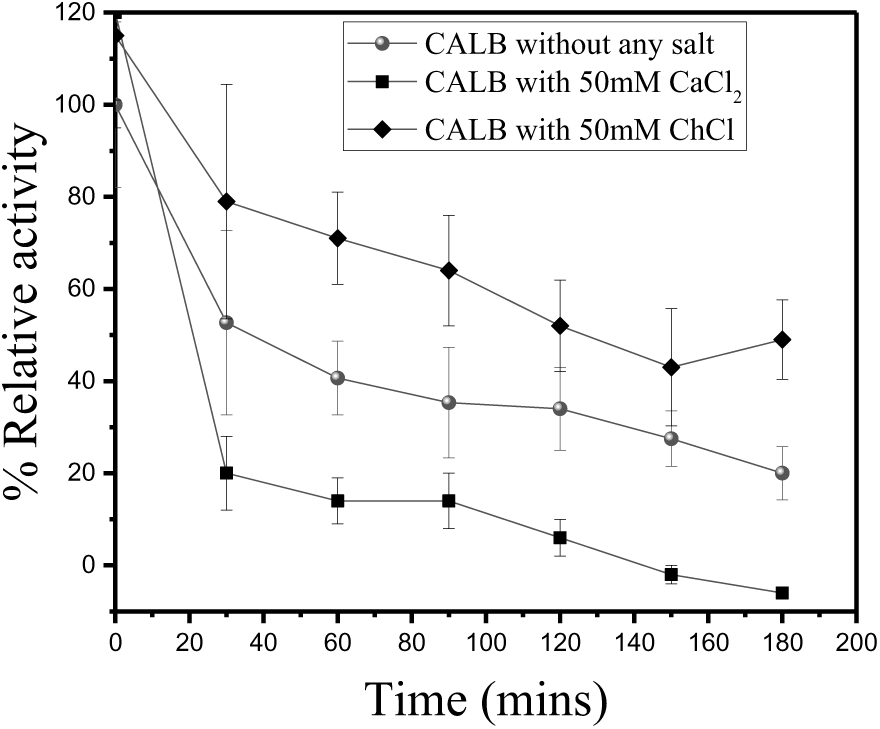
Thermal stability of the CALB enzyme at 50 °C. Enzyme (CALB) without any effector is represented by spheres (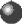), enzyme with 50 mM CaCl_2_ is represented by squares (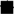) and enzyme with 50mM ChCl is represented by the diamonds (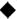).

### 3.6 Surfactant activation

The activation of lipases by surfactant has been reported. In particular, marked activation by cationic surfactants has been shown for a lipase from *Thermomyces lanuginosa*.^28^ Figure 11 shows the effect of cationic and anionic surfactants on CALB. The cationic surfactant (CTAB) showed activation of CALB, by 1.7-fold, at 10 µM, further increasing to 50 and 100 µM, the activity dropped significantly, equivalent to control. Though CALB can be activated by CTAB, the activation was substantially less than that observed for alkylammonium salts (see Figure 4 and Figure 5). When an anionic surfactant, SDS, was tested, no significant activation of CALB was observed over the concentration range from 10 µM to 5mM as shown in Figure 11 (b). Further increasing the SDS concentration to 50mM inhibits the CALB activity probably due to detergent effects. Clearly, the presence of the positive charge on the choline moiety of CTAB was required for activation.

**Figure 11:**
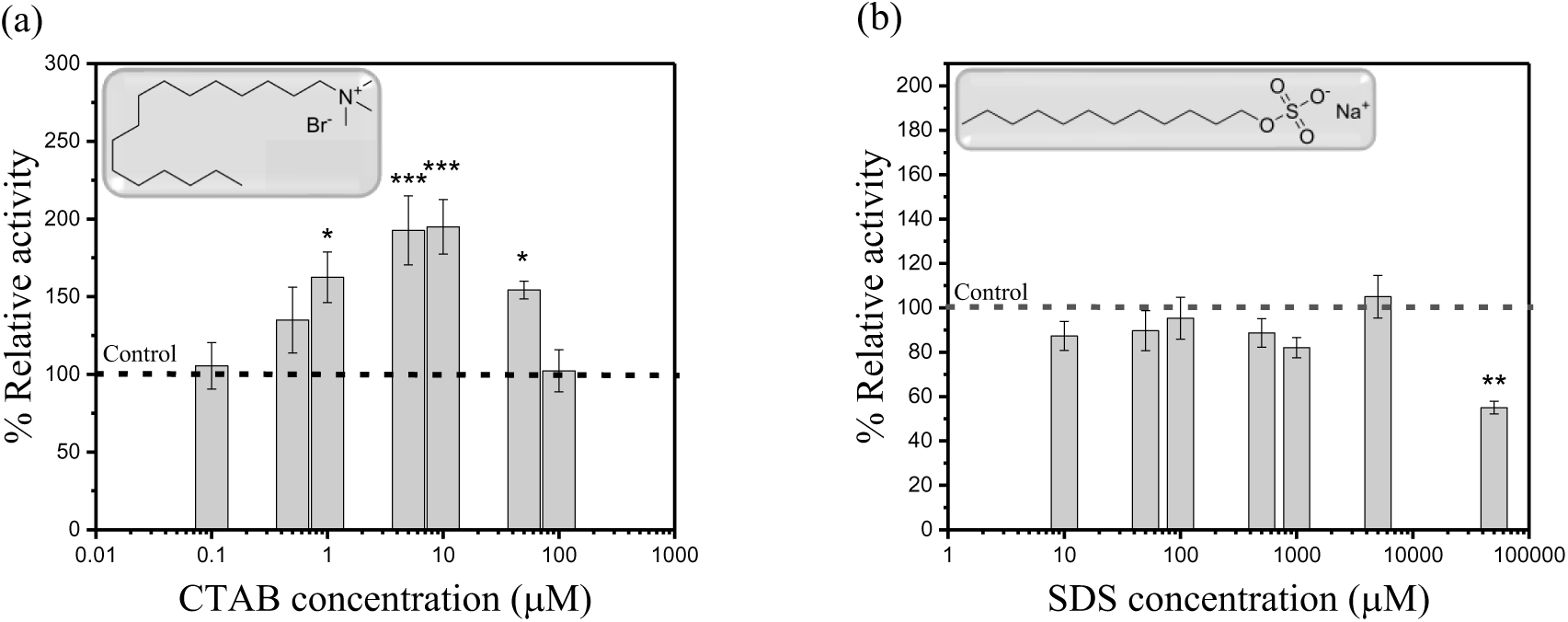
(a) Activation of CALB by the cationic surfactant, CTAB (cetyltrimethylammonium bromide) and (b) effect of the anionic surfactant, SDS (sodium dodecyl sulphate). Activity was measured using the standard assay (see methods above). CTAB from 0.1 to 100 µM and SDS from 10 µM to 10 mM were investigated. The dotted line represents the control. * represents the p value < 0.05 for a confidence interval of 95% and ** represents p value < 0.01 for confidence interval of 99 %.

Taken together, the data above show that, the activation observed on hydrophobic supports, by surfactants and by DES can all be explained by a hydrophobic interaction with a site on the surface of CALB, centered around Asp-145. However, the calcium binding site appears to act independently to cause CALB activation. Ultimately, these sites both act to increase enzyme turnover and must therefore, be mechanistically related. Further studies are needed to fully define the relationship between these two sites.

## 4 General discussion

Enzyme activation in DES has been variously explained as being due to bulk solvent properties, to the formation of a hydrogen bonded lattice that stabilises a specific high activity conformation, to direct interaction with amino acids at the catalytic centre^66^, H-bonding interactions^42^, charge transfer issues etc.^41^ Herein, we show that the mechanism of activation for CALB is, in fact, due to binding of the hydrogen bond acceptor component of DES, choline. Various findings in the literature confirm that others have observed this phenomenon albeit, in most cases, indirectly (see Table 1). Similar effects were observed for Horse Liver Alcohol Dehydrogenase.^41^ These findings are significant in proving that activation is not simply a bulk solvent effect but rather due to the interaction of a component of the DES with the enzyme. What is remarkable is that this finding adds to a growing number of examples of choline activation of enzymes.^36,48,67–69^ Cumulatively, they suggest that such activation is widespread.

Herein, we show that choline is, indeed, an activator of CALB and that the activation is due to the presence of the quaternary ammonium moiety of choline. This allowed us to discount the hydrogen bonding theory or the formation of a supramolecular net as activation mechanisms. Furthermore, we show that the hydrophobic alkyl corona surrounding the nitrogen ion greatly influences the degree of activation. Surprisingly, quaternary nitrogen compounds bearing long alkyl chains can provide extraordinary activation of CALB. The most striking observation is, tetraoctylammonium salt showing an ∼ 5-fold activation of CALB at as low as micromolar (µM) levels. This activation is unprecedented and is, to the best of our knowledge, the highest degree of activation for a small molecule, non-specific activator, reported to date, for this or any enzyme. This work establishes these salts as a new class of activator for biocatalysis that are applicable to a wide number of enzymes. Importantly, the mechanism of activation described herein provides an explanation that reconciles the various types of CALB activation. Thus, it shows that the activation seen in DES, the activation upon immobilisation on hydrophobic supports and the activation by in detergents with long alkyl chains can all be traced to a hydrophobic patch on the enzyme surface.

This work provides, for the first time, a soluble activator for mechanistic studies that can be used to optimise activation and concomitant stabilisation. It may also be useful for enzyme stabilisation via immobilisation. It is worth noting that the activation and stabilisation of immobilised enzymes has always been ascribed to their multi-point attachment to the support matrix. However, at least in this case, the stabilisation noted for immobilised CALB may be due to hydrophobic interaction with a region on the enzyme surface alone. It is unlikely that this is the greatest degree of activation achievable for CALB and further optimisation will be possible. Ideally, a crystal structure of the enzyme-bound activator is needed to confirm the binding sites suggested here.

*Metal ion activation:* Prior to this work, the only other activating agent reported that was not likely to be due to a hydrophobic effect was activation by calcium ions. We showed that calcium activated CALB, presumably by binding at a site that straddles two negatively charged residues on the enzyme, as seen with other proteins.^64^ To our surprise, the addition of choline chloride when calcium activation was at its highest level (100mM), further increased CALB activity. This showed that these sites of interaction were independent. It is not surprising that a divalent cation and a monovalent cationic alkylammonium ion do not bind at the same site. However, the fact that maximum rate enhancement with calcium ions could be further increased by choline addition was intriguing. When stability was examined, a divergence between the two compounds, calcium and ChCl, was observed. We found that calcium destabilised CALB while choline, by contrast, stabilised the enzyme. Taken together this mixed activator experiment suggests that two independent sites are capable of CALB activation.

*Mechanism of CALB activation in DES*: This work explains the activation of CALB in DES, through direct binding of alkylammonium salts to a hydrophobic patch surrounding a negatively charged residue on the enzyme surface. It is possible that the DES-mediated activation of CALB could be due to cholinium components of DES binding to a specific part of the enzyme and that this binding site consists of a negative charge surrounded by a hydrophobic patch (see Figure 12). Exploiting this binding interaction may be a means to activate and stabilise a wide range of enzymes.

**Figure 12:**
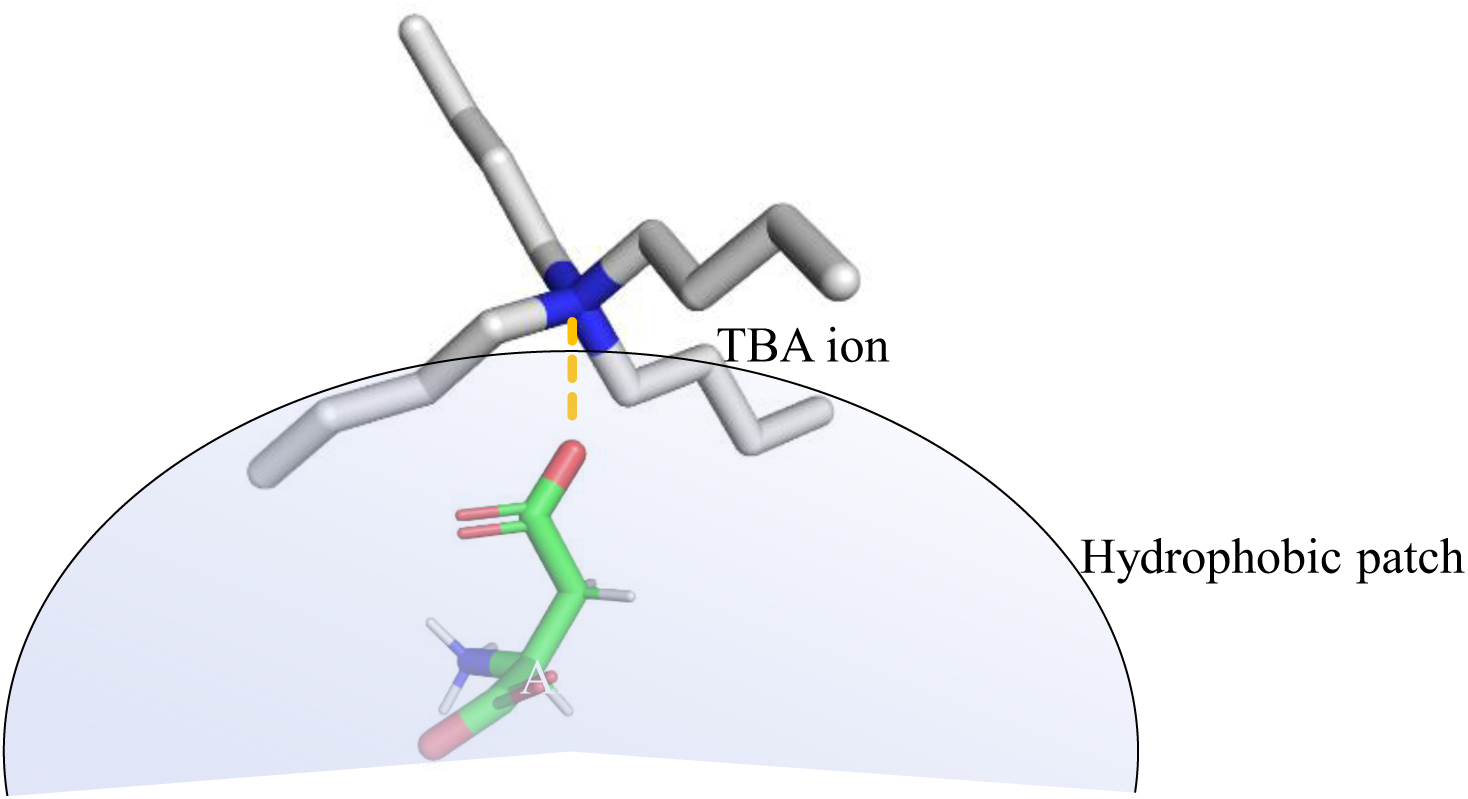
Schematic illustration of tetrabutylammonium (TBA) ion binding to the hydrophobic region on the surface of CALB centered around Asp-145. The shaded area represents the hydrophobic patch on the surface of the enzyme. The electrostatic interaction between the negatively charged aspartate and the positively charged quaternary nitrogen is indicated by a yellow dotted line.

*Novel class of enzyme activators:* The quaternary ammonium salts represent a new class of soluble enzyme activators unlike any previously used activators. Molecules that are capable of activating more than one enzyme (outside of specific peptides binding to allosteric sites) are unknown. The TAAs represent a new class of such compounds with the unique ability to activate a wide range of enzymes.

As has been pointed out by several workers, the search for enzyme activators as therapeutics has lagged the use of inhibitors for a host of reasons.^70^ The fact that this novel class of activators has broad applicability suggests a new binding modality. Thus, the model is that the charge interaction at the centre of the quaternary salt orientates the activator on the enzyme surface “landing site”. The accompanying hydrophobic interactions serve to enhance binding (see Figure 12).

*Paradigm shift:*

If activation in DES is not due to a bulk solvent effect such as hydrogen bonding or charge interactions with the enzyme surface, then the view of DES changes considerably. Thus, it is clear DES are not imbued with unique properties but simply contain components that are activatory (e.g. choline) or inhibitory (e.g. glycerol).

*Understanding activation?* The fundamental processes underlying activation are not yet clear. The observation that so many enzymes become activated in DES must relate in some way to their physiological role *in vivo*. Thus, it is reasonable to deduce that such activation has a physiological counterpart. Is there a process *in vivo* that similarly activates enzymes? Is it possible that isolated enzymes might deviate significantly in terms of activity from those in dilute aqueous solution? Two recent strands of research are useful to consider regarding the enzyme’s behaviour in DES; the intracellular environment and the formation of transient enzyme associations (see below).

*Intracellular environment*: Enzymes have evolved in the environment of the cell where they do not encounter the high solute concentrations of specific components present in DES. Intracellular concentrations of choline, for example, are thought to be in the region of 5-10 µM.^71^ In DES, choline concentrations can be in the millimolar range and above, depending on the level of its inclusion in reaction mixtures. Until DES came to prominence as reaction media, the use of enzymes in the presence of high solutes had not been well-studied. The behaviour of enzymes in very high solute concentrations might be expected to be unusual and it is possible to imagine all kinds of aberrant salt effects, hydrogen bonding interactions etc attributable to this environment. In this regard, it is worth noting that some researchers have pointed out that the intracellular milieu of a cell may be closer in character to a DES than is generally recognised. NMR-based metabolomic studies revealed that all cells contain considerable amounts of certain small molecules such as choline, various sugars, some amino acids and some organic acids (malic acid, citric acid, lactic acid and succinic acid) at a level too high to be considered simply as reaction intermediates in cellular metabolism. Rather, they may be responsible for the establishment of an intracellular DES-like environment. The authors argue that the metabolic activity of the cell takes place in what is effectively a eutectic mixture. Historically, studies of enzymes have been carried out in dilute, buffered, aqueous solutions which does not fully reflect their native environment. More recently, an exploration of osmolytes (compounds produced in cells as a response to adverse conditions) showed that they can give rise to complex eutectic mixtures.^72^ Thus, DES activation may simply reflect the fact that many enzymes evolved to operate optimally in a DES-like environment.

*Aggregation phenomena*: Given that CALB is activated by hydrophobic interactions, it is conceivable that hydrophobic patches on adjacent CALB monomers could bind each other to create a more active dimer or higher order multimeric protein. Indeed, changes in the state of aggregation of enzymes are becoming more widely appreciated and there is evidence that this process can occur in response to various effectors.^73^ The formation of catalytically enhanced dimeric species is known for lipases.^74^ Activation by TAA salts could act to intercept this aggregation. In this model, the alkylammonium salts act to occupy the “dimerization*”* site causing the observed activation - essentially preventing enzyme association. This is, indeed, possible and may be a likely explanation for this “choline effect”. Moreover, the formation of bimolecular condensates is known to accelerate enzyme reactions.^75^ Arguing against this explanation is the observation that CALB, reportedly, does not exhibit dimerization^76^ although a self-activation phenomenon has been reported for CALB with the involvement of a “lid holder” in the open chain form of CALB.^77^

In a broader sense, to the best of our knowledge, choline is the first physiological compound known to activate CALB. Clearly, choline-containing lipids might interact with this binding site where affinity, due to proximity, might be locally enhanced and choline containing lipids are present in *Candida*. Of course, ChCl is one of several quaternary nitrogen compounds present in cells and this study cannot discount the possibility that many cell components (phosphatidyl choline, lysophosphatidyl choline, carnitine, trimethyl amine oxide *etc*.) bearing this moiety could also activate CALB.

The quaternary nitrogen compounds studied herein are essentially multi-directional hydrophobic patches solubilised by the permanent charge on the nitrogen atom. It is possible to envisage their adherence to sites on the enzyme involved in the formation of aggregates or condensates, effectively disrupting the aggregation process. Indeed, recent studies suggest that the formation of condensates is widespread and is governed by hydrophobic interactions.^78^ It is also well-established that for many enzymes, including lipases, that condensate formation gives rise to activation and stabilisation. Indeed, the proximity of enzymes in Cross-Linked Enzyme Aggregates (CLEAs) is well known to stabilise and activate biocatalysts.^79^

If TAA compounds are effectively acting to disaggregate proteins while retaining their activation, then these molecules could find application in disorders of protein aggregation, as therapeutics and probes of mechanism.

*Biomedical applications*: A number of review articles have highlighted the need for consideration of enzyme activation as a viable therapeutic strategy. Thus, activation of enzymes such as Pyruvate Kinase and Glucokinase have been pursued as therapeutic objectives. It is possible to envisage other areas where targeted activation could be pursued. This work shows that the exploration of enzyme activators and activation pathways may be a profitable avenue of future research for therapeutic application.

We are aware that this work did not optimise binding of TAA and that compounds with even greater efficacy may be found using these compounds as starting points. We hope this report will stimulate further research on small molecule enzyme activators as an attractive alternative to mutagenesis, immobilisation or solvent engineering approaches.

## Acknowledgement

This research was funded under a Science Foundation Ireland (SFI) Programme, Frontiers for the Future: grant number 21/FFP-A/9898

## Data availability

All relevant data is provided in the manuscript and supplementary information of this work.

## Supplementary information

There is a linked supplementary information document for this manuscript

## Abbreviations

CALB: *Candida antarctica* Lipase B
ChCl: Choline chloride
CLEA: Cross-Linked Enzyme Aggregate
CTAB: Cetyltrimethylammonium bromide
DES: Deep eutectic solvent
Gly: Glycerol
HBA: Hydrogen bond acceptor
HBD: Hydrogen bond donor
HLADH: Horse liver alcohol dehydrogenase
MTOAC: Methyltrioctylammonium chloride
NADES: Natural Deep Eutectic Solvents
TAA: Tetraalkylammonium
TBAB: Tetrabutylammonium bromide
TBAC: Tetrabutylammonium chloride
TEAB: Tetraethylammonium bromide
THAB: Tetrahexylammonium bromide
TMAB: Tetramethylammonium bromide
TOAB: Tetraoctylammonium bromide
TDAB: Tetrakis(decyl)ammonium bromide

## References

1. Zaks, A. & Klibanov, M. A. Enzymatic Catalysis in Organic Media at 100 °C. Science (1979) 224, (1984).

2. Sheldon, R. A. & Brady, D. Streamlining design, engineering, and applications of enzymes for sustainable biocatalysis. ACS Sustainable Chemistry and Engineering vol. 9 8032–8052 Preprint at 10.1021/acssuschemeng.1c01742 (2021).

3. Kissman, E. N. et al. Expanding chemistry through in vitro and in vivo biocatalysis. Nature vol. 631 37–48 Preprint at 10.1038/s41586-024-07506-w (2024).

4. Paul, C. et al. Enzyme engineering for biocatalysis. Molecular Catalysis 555, (2024).

5. Zhou, D., Chen, X., Li, G., Zhao, M. & Li, D. Effect of deep eutectic solvents on activity, stability, and selectivity of enzymes: Novel insights and further prospects. International Journal of Biological Macromolecules vol. 284 Preprint at 10.1016/j.ijbiomac.2024.138148 (2025).

6. Giri, P., Pagar, A. D., Patil, M. D. & Yun, H. Chemical modification of enzymes to improve biocatalytic performance. Biotechnology Advances vol. 53 Preprint at 10.1016/j.biotechadv.2021.107868 (2021).

7. Sheldon, R. A. & Pereira, P. C. Biocatalysis engineering: The big picture. Chemical Society Reviews vol. 46 2678–2691 Preprint at 10.1039/c6cs00854b (2017).

8. Nian, B. & Li, X. Can deep eutectic solvents be the best alternatives to ionic liquids and organic solvents: A perspective in enzyme catalytic reactions. International Journal of Biological Macromolecules vol. 217 255–269 Preprint at 10.1016/j.ijbiomac.2022.07.044 (2022).

9. Baby, E. K. et al. Influence of deep eutectic solvents on redox biocatalysis involving alcohol dehydrogenases. Heliyon vol. 10 Preprint at 10.1016/j.heliyon.2024.e32550 (2024).

10. Fries, R. W., Bohlken, D. P., Blakley, R. T. & Plapp, B. V. Activation of Liver Alcohol Dehydrogenases by Imidoesters Generated in Solution”! DUBLIN CITY UNIV on 14, 5233–5238 (1975).

11. Fang, Y. et al. N-terminal lid swapping contributes to the substrate specificity and activity of thermophilic lipase TrLipE. Front Microbiol 14, (2023).

12. Niu, K. et al. Artificial trimerization of β-glucosidase for enhanced thermostability and activity via computational redesign. Int J Biol Macromol 286, (2025).

13. Wang, Z. et al. Computational Redesign of the Substrate Binding Pocket of Glutamate Dehydrogenase for Efficient Synthesis of Noncanonical l-Amino Acids. ACS Catal 12, 13619–13629 (2022).

14. Pätzold, M. et al. Deep Eutectic Solvents as Efficient Solvents in Biocatalysis. Trends in Biotechnology vol. 37 943–959 Preprint at 10.1016/j.tibtech.2019.03.007 (2019).

15. Sheldon, R. A. & Woodley, J. M. Role of Biocatalysis in Sustainable Chemistry. Chemical Reviews vol. 118 801–838 Preprint at 10.1021/acs.chemrev.7b00203 (2018).

16. Sakhuja, D., et al. Cost-effective production of biocatalysts using inexpensive plant biomass: a review. 3 Biotech vol. 11 Preprint at 10.1007/s13205-021-02847-z (2021).

17. Safdar, A., Ismail, F. & Imran, M. Biodegradation of synthetic plastics by the extracellular lipase of Aspergillus niger. Environmental Advances 17, (2024).

18. Magalhães, R. P., Cunha, J. M. & Sousa, S. F. Perspectives on the role of enzymatic biocatalysis for the degradation of plastic pet. International Journal of Molecular Sciences vol. 22 Preprint at 10.3390/ijms222011257 (2021).

19. Freije García, F. & García Liñares, G. Use of Lipases as a Sustainable and Efficient Method for the Synthesis and Degradation of Polymers. Journal of Polymers and the Environment vol. 32 2484–2516 Preprint at 10.1007/s10924-023-03118-z (2024).

20. Chandra, P., Enespa, Singh, R. & Arora, P. K. Microbial lipases and their industrial applications: A comprehensive review. Microbial Cell Factories vol. 19 Preprint at 10.1186/s12934-020-01428-8 (2020).

21. Waggett, A. & Pfaendtner, J. Hydrophobic Residues Promote Interfacial Activation of Candida rugosa Lipase: A Study of Rotational Dynamics. Langmuir (2024) doi:10.1021/acs.langmuir.4c02174.

22. Reis, P., Holmberg, K., Watzke, H., Leser, M. E. & Miller, R. Lipases at interfaces: A review. Advances in Colloid and Interface Science vols 147–148 237–250 Preprint at 10.1016/j.cis.2008.06.001 (2009).

23. Zisis, T. et al. Interfacial Activation of Candida antarctica Lipase B: Combined Evidence from Experiment and Simulation. Biochemistry 54, 5969–5979 (2015).

24. Stauch, B., Fisher, S. J. & Cianci, M. Open and closed states of Candida Antarctica lipase B: Protonation and the mechanism of interfacial activation. J Lipid Res 56, 2348–2358 (2015).

25. Ortiz, C. et al. Novozym 435: The ‘perfect’ lipase immobilized biocatalyst? Catalysis Science and Technology vol. 9 2380–2420 Preprint at 10.1039/c9cy00415g (2019).

26. Pazol, J., Weiss, T. M., Martínez, C. D., Quesada, O. & Nicolau, E. The influence of calcium ions (Ca2+) on the enzymatic hydrolysis of lipopolysaccharide aggregates to liberate free fatty acids (FFA) in aqueous solution. JCIS Open 7, (2022).

27. Cui, J., Zhao, Y., Liu, R., Zhong, C. & Jia, S. Surfactant-activated lipase hybrid nanoflowers with enhanced enzymatic performance. Sci Rep 6, (2016).

28. Moreno-Perez, S., Ghattas, N., Filice, M., Guisan, J. M. & Fernandez-Lorente, G. Dramatic hyperactivation of lipase of Thermomyces lanuginosa by a cationic surfactant: Fixation of the hyperactivated form by adsorption on sulfopropyl-sepharose. J Mol Catal B Enzym 122, 199–203 (2015).

29. Siódmiak, T., Dulęba, J., Haraldsson, G. G., Siódmiak, J. & Marszałł, M. P. The Studies of Sepharose-Immobilized Lipases: Combining Techniques for the Enhancement of Activity and Thermal Stability. Catalysts 13, (2023).

30. Li, C. et al. Self-assembly of activated lipase hybrid nanoflowers with superior activity and enhanced stability. Biochem Eng J 158, (2020).

31. Freitas, L., Bueno, T., Perez, V. H., Santos, J. C. & De Castro, H. F. Enzymatic hydrolysis of soybean oil using lipase from different sources to yield concentrated of polyunsaturated fatty acids. World J Microbiol Biotechnol 23, 1725–1731 (2007).

32. Hertadi, R. & Widhyastuti, H. Effect of Ca2+ Ion to the Activity and Stability of Lipase Isolated from Chromohalobacter japonicus BK-AB18. Procedia Chem 16, 306–313 (2015).

33. Kukreja, V. & Bera, M. B. Lipase from Pseudomonas Aeruginosa MTCC 2488: Partial Purification, Characterization and Calcium Dependent Thermostability. Indian Journal of Biotechnology vol. 4 (2005).

34. Gomi, K., Ota, Y. & Minoda, Y. Role of lipase activators produced by saccharomycopsis lipolytica and calcium ion in its lipase reaction. Agric Biol Chem 50, 2531–2536 (1986).

35. Dimitrijević, A. et al. One-step, inexpensive high yield strategy for Candida antarctica lipase A isolation using hydroxyapatite. Bioresour Technol 107, 358–362 (2012).

36. Huang, Z. L., Wu, B. P., Wen, Q., Yang, T. X. & Yang, Z. Deep eutectic solvents can be viable enzyme activators and stabilizers. Journal of Chemical Technology and Biotechnology 89, 1975–1981 (2014).

37. Logarušić, M. et al. Harnessing the potential of deep eutectic solvents in biocatalysis: design strategies using CO2 to formate reduction as a case study. Front Chem 12, (2024).

38. Arnodo, D., Maffeis, E., Marra, F., Nejrotti, S. & Prandi, C. Combination of Enzymes and Deep Eutectic Solvents as Powerful Toolbox for Organic Synthesis. Molecules vol. 28 Preprint at 10.3390/molecules28020516 (2023).

39. Abranches, D. O. & Coutinho, J. A. P. Everything You Wanted to Know about Deep Eutectic Solvents but Were Afraid to Be Told. (2023) doi:10.1146/annurev-chembioeng.

40. Zhang, N. et al. Redox Biocatalysis in Lidocaine-Based Hydrophobic Deep Eutectic Solvents: Non-Conventional Media Outperform Aqueous Conditions. ChemSusChem (2024) doi:10.1002/cssc.202402075.

41. Baby, E. K. et al. Novel alcohol dehydrogenase activation by the choline component of Deep Eutectic Solvents. J Mol Liq 423, (2025).

42. Nian, B., Cao, C. & Liu, Y. How Candida antarctica lipase B can be activated in natural deep eutectic solvents: experimental and molecular dynamics studies. Journal of Chemical Technology and Biotechnology 95, 86–93 (2020).

43. Tan, Y., Henehan, G. T., Kinsella, G. K. & Ryan, B. J. Cutinase from Amycolatopsis mediterannei: Marked activation and stabilisation in Deep Eutectic Solvents. Bioresour Technol Rep 16, (2021).

44. Zheng, X., et al. Deep eutectic solvent as an additive to improve enzymatic hydrolysis of polyethylene terephthalate (PET). Preprint, (2024).

45. Juneidi, I., Hayyan, M., Hashim, M. A. & Hayyan, A. Pure and aqueous deep eutectic solvents for a lipase-catalysed hydrolysis reaction. Biochem Eng J 117, 129–138 (2017).

46. Rozas, S., Benito, C., Alcalde, R., Atilhan, M. & Aparicio, S. Insights on the water effect on deep eutectic solvents properties and structuring: The archetypical case of choline chloride + ethylene glycol. J Mol Liq 344, (2021).

47 . De María, P. D., Guajardo, N. & Kara, S. Enzyme catalysis: In DES, with DES, and in the presence of DES. in Deep Eutectic Solvents: Synthesis, Properties, and Applications 257–271 (wiley, 2019). doi:10.1002/9783527818488.ch13.

48. Chan, K. K. et al. Enhanced activity of Candida antarctica lipase B in cholinium aminoate ionic liquids: a combined experimental and computational analysis. J Biomol Struct Dyn 42, (2023).

49. Chan, K. K. et al. Investigation of laccase activity in cholinium-based ionic liquids using experimental and molecular dynamics techniques. Int J Biol Macromol 277, (2024).

50. Xu, W. J. et al. Improving Β-glucosidase biocatalysis with deep eutectic solvents based on choline chloride. Biochem Eng J 138, 37–46 (2018).

51. Wu, B. P., Wen, Q., Xu, H. & Yang, Z. Insights into the impact of deep eutectic solvents on horseradish peroxidase: Activity, stability and structure. J Mol Catal B Enzym 101, 101–107 (2014).

52. Zhao, H., Baker, G. A. & Holmes, S. New eutectic ionic liquids for lipase activation and enzymatic preparation of biodiesel. Org Biomol Chem 9, 1908–1916 (2011).

53. Deive, F. J. et al. On the hunt for truly biocompatible ionic liquids for lipase-catalyzed reactions. RSC Adv 5, 3386–3389 (2015).

54. Fernandez-Lafuente, R., Armisén, P., Sabuquillo, P., Fernández-Lorente, G. & Guisán, J. M. Immobilization of Lipases by Selective Adsorption on Hydrophobic Supports. Chemistry and Physics of Lipids vol. 93 (1998).

55. de Andrades, D. et al. Effect of the support alkyl chain nature in the functional properties of the immobilized lipases. Enzyme Microb Technol 184, (2025).

56. Armonk. IBM SPSS Statistics for Windows. Preprint at (2024).

57. Schrödinger. The PyMOL Molecular Graphics System. Preprint at (2024).

58. Huey, R., Morris, G. M., Olson, A. J. & Goodsell, D. S. A semiempirical free energy force field with charge-based desolvation. J Comput Chem 28, 1145–1152 (2007).

59. Eberhardt, J., Santos-Martins, D., Tillack, A. F. & Forli, S. AutoDock Vina 1.2.0: New Docking Methods, Expanded Force Field, and Python Bindings. J Chem Inf Model 61, 3891–3898 (2021).

60. Trott, O. & Olson, A. J. AutoDock Vina: Improving the speed and accuracy of docking with a new scoring function, efficient optimization, and multithreading. J Comput Chem 31, 455–461 (2010).

61. BIOVIA Discovery Studio Visualizer. Preprint at (2024).

62. Nian, B., Cao, C. & Liu, Y. Activation and stabilization of Candida antarctica lipase B in choline chloride-glycerol-water binary system via tailoring the hydrogen-bonding interaction. Int J Biol Macromol 136, 1086–1095 (2019).

63. Pardo-Tamayo, J. S., Arteaga-Collazos, S., Domínguez-Hoyos, L. C. & Godoy, C. A. Biocatalysts Based on Immobilized Lipases for the Production of Fatty Acid Ethyl Esters: Enhancement of Activity through Ionic Additives and Ion Exchange Supports. BioTech 12, (2023).

64. Mcphaien, C. A., Strynadka, N. C. J. & James, M. N. G. CALCIUM-BINDING SITES IN PROTEINS: A STRUCTURAL PERSPECTIVE. (1991).

65. Kumar, A. & Venkatesu, P. Overview of the stability of α-chymotrypsin in different solvent media. Chemical Reviews vol. 112 4283–4307 Preprint at 10.1021/cr2003773 (2012).

66. Toledo, M. L. et al. Laccase Activation in Deep Eutectic Solvents. ACS Sustain Chem Eng 7, 11806–11814 (2019).

67. Nascimento, P. A. M., Picheli, F. P., Lopes, A. M., Pereira, J. F. B. & Santos-Ebinuma, V. C. Effects of cholinium-based ionic liquids on Aspergillus niger lipase: Stabilizers or inhibitors. Biotechnol Prog 35, (2019).

68. Hoppe, J. et al. An effect of choline lactate based low transition temperature mixtures on the lipase catalytic properties. Colloids Surf B Biointerfaces 216, (2022).

69. Zhao, H., Baker, G. A. & Holmes, S. Protease activation in glycerol-based deep eutectic solvents. J Mol Catal B Enzym 72, 163–167 (2011).

70. Dow, L. F. et al. The evolution of small molecule enzyme activators. RSC Medicinal Chemistry vol. 14 2206–2230 Preprint at 10.1039/d3md00399j (2023).

71. Sarter, M. & Parikh, V. Choline transporters, cholinergic transmission and cognition. Nature Reviews Neuroscience vol. 6 48–56 Preprint at 10.1038/nrn1588 (2005).

72. Kim, S. H. et al. Effect of deep eutectic solvent mixtures on lipase activity and stability. J Mol Catal B Enzym 128, 65–72 (2016).

73. Gentile, K. et al. Enzyme aggregation and fragmentation induced by catalysis relevant species. Physical Chemistry Chemical Physics 23, 20709–20717 (2021).

74. Fernández-Lorente, G. et al. Self-assembly of Pseudomonas fluorescens lipase into bimolecular aggregates dramatically affects functional properties. Biotechnol Bioeng 82, 232–237 (2003).

75. Dindo, M. et al. Enzymes can activate and mobilize the cytoplasmic environment across scales. Preprint at 10.1101/2025.01.28.635259 (2025).

76. Palomo, J. M. et al. General Trend of Lipase to Self-Assemble Giving Bimolecular Aggregates Greatly Modifies the Enzyme Functionality. (2003) doi:10.1021/bm025729.

77. Luan, B. & Zhou, R. A novel self-activation mechanism of: Candida antarctica lipase B. Physical Chemistry Chemical Physics 19, 15709–15714 (2017).

78. Krainer, G. et al. Reentrant liquid condensate phase of proteins is stabilized by hydrophobic and non-ionic interactions. Nat Commun 12, (2021).

79. Velasco-Lozano, S., López-Gallego, F., Mateos-Díaz, J. C. & Favela-Torres, E. Cross-linked enzyme aggregates (CLEA) in enzyme improvement – a review. Biocatalysis 1, (2016).

