## Supporting information for "Extraordinary activation of CALB by alkylammonium ions: a new paradigm for activity enhancement of enzymes"

### Effect of sodium chloride on CALB activity

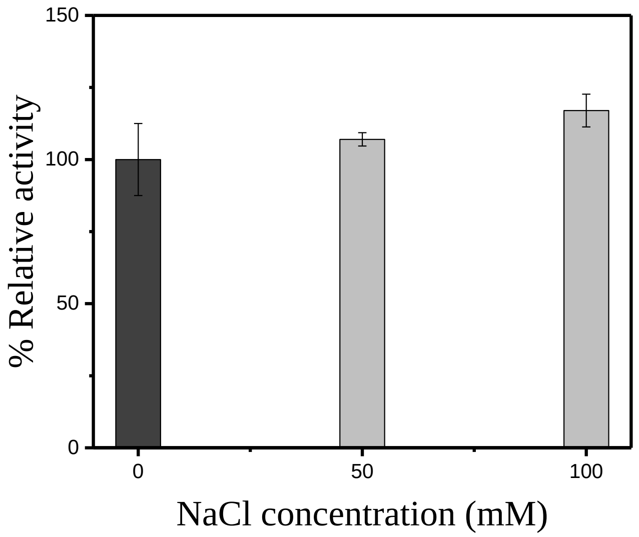


Figure S1: CALB in the presence of 50 and 100 mM NaCl showed no significant activation compared to the control enzyme

**Effect of bromide salt of tetrabutylammonium ion on CALB activity**

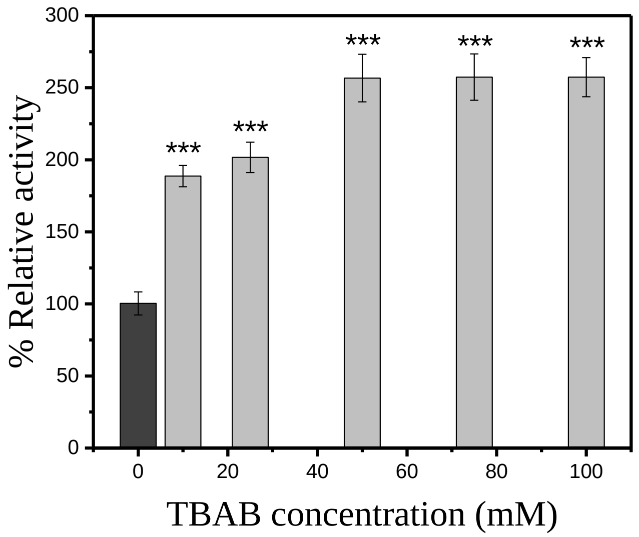


Figure S2: CALB activation in the presence of tetrabutylammonium bromide. The activation by the bromide salt is similar to the activation by chloride salt.

**Effect of different divalent metal ions on CALB activity**

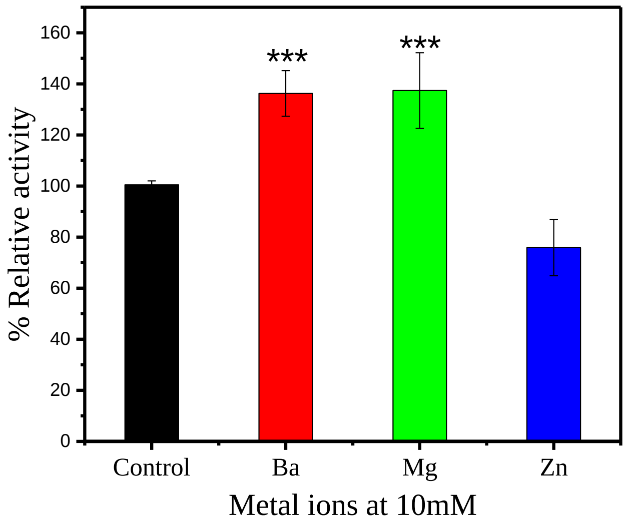


Figure S3: Effect of metal ions at 10mM concentration on CALB activity. $Ba^{2+}$ and $Mg^{2+}$ showed significant activation, $Zn^{2+}$ shows inhibition of CALB

### Effect of combining choline chloride and tetrabutylammonium bromide

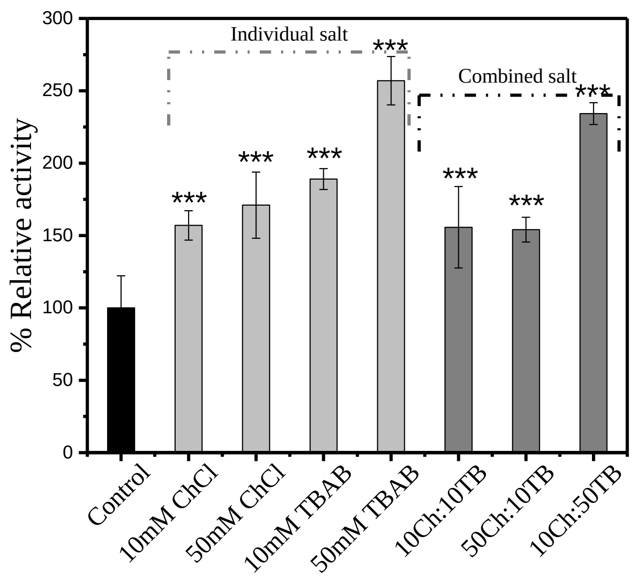


Figure S4: The combined addition of ChCl and TBAB showed competitive behaviour, where the resultant CALB activation is lesser than that achieved with individual salts.

Table S1 Accessible surface areas of negatively charged amino acids on the surface of chain B of CALB.

| **Amino Acid** | **Accessible Surface Area (Å)** |
|---|---|
| ASP 6 | 47.00 |
| ASP 17 | 50.59 |
| ASP 49 | 78.90 |
| ASP 75 | 21.03 |
| GLU 81 | 1.83 |
| ASP 126 | 26.55 |
| ASP 134 | 2.77 |
| ASP 145 | 30.43 |
| ASP 187 | 2.03 |
| GLU 188 | 79.94 |
| ASP 223 | 60.82 |
| ASP 252 | 53.44 |
| ASP 257 | 32.52 |
| ASP 265 | 80.11 |
| GLU 269 | 128.75 |
| GLU 294 | 8.67 |
| ASP 296 | 50.50 |

Table S2 Accessible surface areas of hydrophobic amino acids surrounding Asp-145 on the surface of chain B of CALB.

| **Amino Acid** | **Accessible Surface Area (Å)** |
|---|---|
| THR 138 | 20.48 |
| VAL 139 | 87.03 |
| LEU 140 | 72.40 |
| ALA 141 | 82.34 |
| GLY 142 | 42.47 |
| PRO 143 | 135.46 |
| LEU 144 | 71.81 |
| ASP 145 | 30.43 |
| ALA 146 | 68.49 |
| LEU 147 | 151.72 |
| ALA 148 | 42.90 |
| VAL 149 | 71.20 |
| SER 150 | 9.30 |
